# An Oncometabolite Isomer Rapidly Induces A Pathophysiological Protein Modification

**DOI:** 10.1101/2020.01.11.902973

**Authors:** Sarah E. Bergholtz, Chloe A. Briney, Susana S. Najera, Minervo Perez, W. Marston Linehan, Jordan L. Meier

## Abstract

Metabolites regulate protein function via covalent and non-covalent interactions. However, manipulating these interactions in living cells remains a major challenge. Here we report a chemical strategy for inducing cysteine S-succination, a non-enzymatic posttranslational modification derived from the oncometabolite fumarate. Using a combination of antibody-based detection and kinetic assays we benchmark the in vitro and cellular reactivity of two novel S-succination “agonists,” maleate and 2-bromosuccinate. Cellular assays reveal maleate to be a more potent and less toxic inducer of S-succination which can activate KEAP1-NRF2 signaling in living cells. By enabling the cellular reconstitution of an oncometabolite-protein interaction with physiochemical accuracy and minimal toxicity, this study provides a methodological basis for better understanding the signaling role of metabolites in disease.

---

An emerging paradigm in cancer biology is that metabolism may function as an epigenetic signal in and of itself.^1-2^ A prototypical example of this phenomenon occurs in the genetic cancer syndrome hereditary leiomyomatosis and renal cell carcinoma (HLRCC). In this disorder mutations in gene encoding the TCA cycle enzyme fumarate hydratase (FH) cause fumarate to accumulate to millimolar levels.^3^ These aberrant levels of fumarate are associated with chromatin hypermethylation and dysregulated gene expression;^4-5^ however, the molecular mechanisms by which this simple organic metabolite drives such profound changes in epigenetic signaling are not completely understood. Fumarate is unique amongst oncometabolites in that it has two physically distinct mechanisms by which it may alter posttranslational modification (PTM) mediated signaling First, fumarate can serve as a competitive inhibitor of 2-ketoglutarate (KG) utilizing enzymes, a mechanism it shares with other oncometabolites such as succinate and 2-hydroxyglutarate.^6-7^ Second, fumarate can react covalently with proteins to form the non-enzymatic modification cysteine S-succination (Fig. 1a).^8^ This non-enzymatic PTM occurs exclusively in HLRCC and a few other pathophysiological contexts.^9-11^ While the targets and stoichiometries of protein

**Figure 1.**
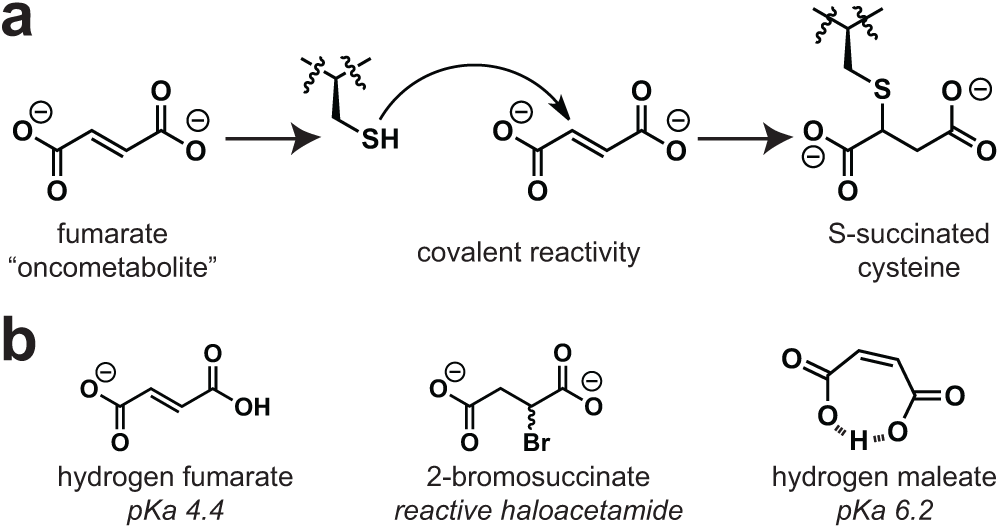
(a) Non-enzymatic modification of cysteines by fumarate causes protein S-succination. (b) Structures of endogenous and synthetic inducers of cellular S-succination.

S-succination are incompletely defined, at least one consequence of this PTM appears to be stabilization of the anti-oxidant transcription factor NRF2 due to covalent S-succination of its regulatory E3 ligase KEAP1.^12-13^

Recently we reported a chemoproteomic approach for characterizing protein S-succination in patient-derived HLRCC cell lines.^14^ Investigation of the local sequence preferences of FH-sensitive cysteine residues and kinetic analyses led to the surprising discovery that fumarate itself appears to be chemically inert towards thiol addition, only becoming reactive upon protonation to hydrogen and dihydrogen fumarate.^15^ This leads to a paradoxical influence of pH on S-succination. However, fumarate’s low reactivity raises a technical challenge: it is very difficult to induce cysteine S-succination in cells. Most studies that have sought to explore fumarate’s reactivity using exogenous S-succination reagents have used dimethyl fumarate (DMF), an ester derivative that implements a physiochemically distinct PTM and is at least ∼100-fold more reactive than endogenous fumarate.^16-17^ Methods to temporally control cysteine S-succination have the potential to decouple fumarate’s covalent and non-covalent epigenetic mechanisms, and provide new insights into the role of this oncometabolite in biology and disease.

To develop a reagent for manipulating oncometabolite-associated S-succination, we envisioned two straightforward modifications to fumarate’s structure. First, given our previous findings, we considered whether fumarate analogues with increased susceptibility to protonation may show coordinately increased cysteine reactivity. This led us to explore maleate, a cis-isomer of fumarate (Fig. 1b). Unlike fumarate, maleate has the unique ability to form an internal hydrogen bond upon protonation, leading to a substantial change in its pKa (maleic acid, pKa_2_ = 6.2; fumaric acid, pKa_2_ = 4.5).^18^ When present at equal concentrations in buffered solution, levels of electrophilic hydrogen maleate are over an order of magnitude higher than hydrogen fumarate, which would be expected to cause a concomitant increase in cysteine S-succination. Second, we hypothesized the intrinsic electrophilicity of fumarate could be increased by replacing the α,β-unsaturated Michael acceptor with a bromoacetamide (Fig. 1b). The cognate analogue, 2-bromosuccinate, has the potential to either label cysteines and form the desired PTM, or undergo an elimination reaction to form electrophilically inert fumarate. Importantly, α-carboxyhaloacetamides have shown previous utility as cysteine labeling reagents in a number of settings.^19-20 21^

As an initial test of the ability of these oncometabolite analogues to induce S-succination, we incubated proteomes with fumarate, maleate, and 2-bromosuccinate and assessed covalent protein labeling using a recently developed anti-cysteine-S-succination antibody (Fig 2). Exposure of proteomes to maleate and 2-bromosuccinate led to protein S-succination that was detectable at concentrations as low as 0.5-1 mM (Fig. 2b). In contrast, exposure of proteomes to fumarate at these low concentrations did not lead to detectable protein labeling, consistent with our previous studies. Comparison of the reagents revealed maleate provided qualitatively greater protein labeling than 2-bromosuccinate when applied at an equivalent concentration (Fig. S1). Both 2-bromosuccinate and maleate hyper-S-succinated proteomes in a time-dependent manner, with signal intensities increasing over 24 hours (Fig. 2c). This suggests that both of these reagents are stable in solution over prolonged time periods in celllysates. These results establish 2-bromosuccinate and maleate as novel synthetic reagents capable of inducing oncometabolite-associated PTM cysteine-S-succination.

**Figure 2.**
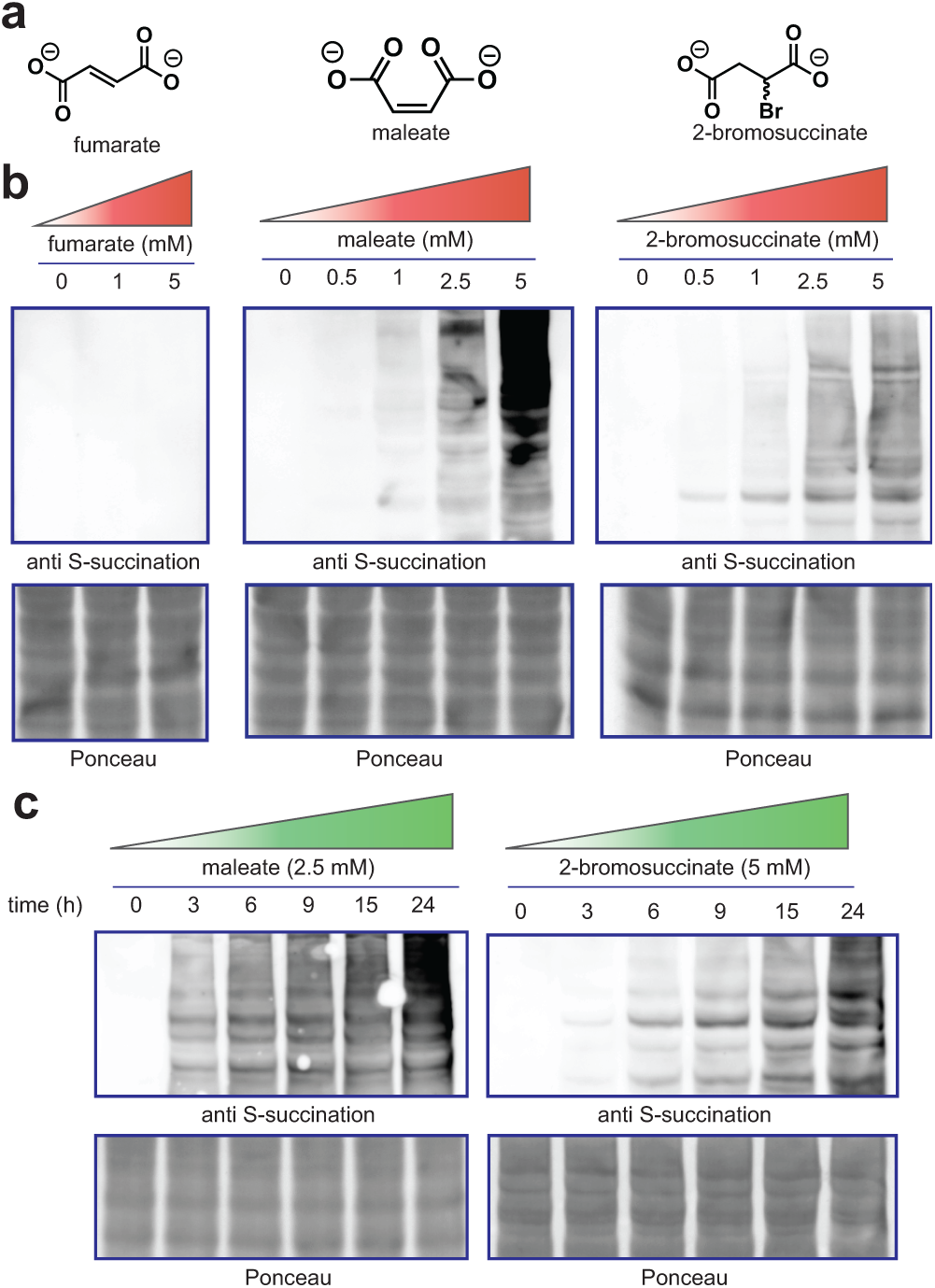
(a) Structure of non-enzymatic S-succination reagents. (b) Dose-dependent effects of fumarate, maleate, and 2-bromosuccinate on protein S-succination in cell lysates (t = 15 h). (c) Time-dependent S-succination of proteomes upon addition of maleate or 2-bromosuccinate.

Next, we aimed to quantitatively benchmark the reactivity of our chemical S-succination reagents. For these experiments, we incubated electrophiles (fumarate, maleate, or 2-bromosuccinate) with a model thiol (thiophenol, pKa 6.6) and assessed product formation via quantitative UV-RP-HPLC (Fig. 3).^14^ Consistent with immunoblotting, these studies revealed that maleate S-succinates thiols much faster (∼30x) than fumarate at neutral pH (Table 1). Examining the influence of pH on thiol modification we verified that fumarate’s reactivity is increased under more acidic conditions. In contrast, maleate and 2-bromosuccinate behave more like a conventional electrophile, with increased reaction rate in neutral or mildly alkali solutions (Table 1). This suggests that unlike hydrogen fumarate (pKa 4.5), sufficient concentrations of hydrogen maleate (pKa 6.2) are present at neutral pH to enable thiol reactivity, the rate of which declines upon acidification due to reduced abundance of thiophenolate nucleophile.

**Table 1.** S-succination reaction kinetics as a function of electrophile and pH.

|  | pH | fumarate | maleate | 2-Br-succinate |
| --- | --- | --- | --- | --- |
| Rate<br>(mM<sup>-1</sup> min<sup>-1</sup>) | 5 | (1.1 ± 0.6) × 10<sup>-4</sup> | (1.6 ± 0.78) × 10<sup>-4</sup> | (0.08 ± 0.01) × 10<sup>-4</sup> |
|  | 6 | (0.11 ± 0.03) × 10<sup>-4</sup> | (2.9 ± 0.66) × 10<sup>-4</sup> | (0.99 ± 0.23) × 10<sup>-4</sup> |
|  | 7 | (0.06 ± 0.01) × 10<sup>-4</sup> | (3.9 ± 0.57) × 10<sup>-4</sup> | (4.2 ± 0.59) × 10<sup>-4</sup> |

**Figure 3.**
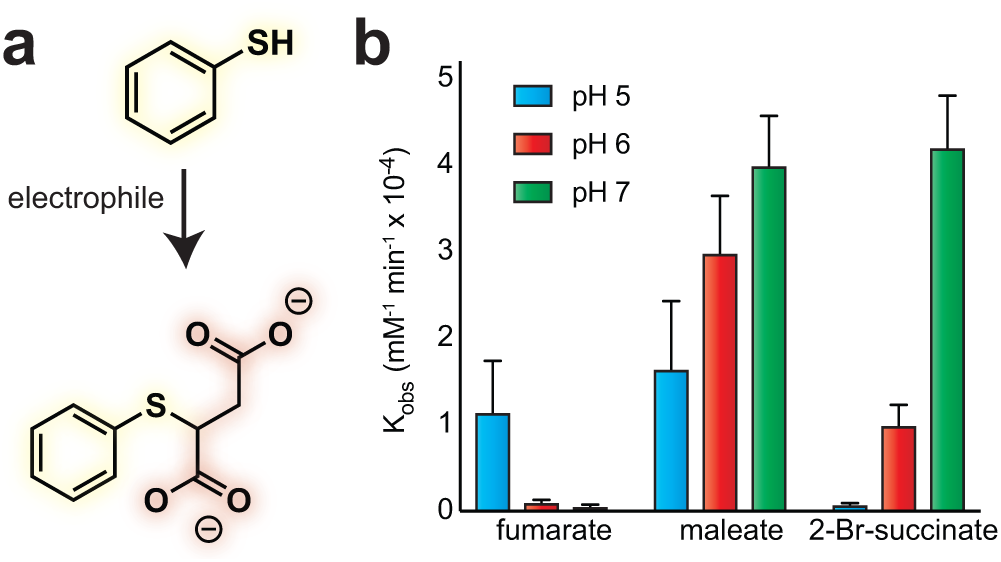
(a) Model reaction to assess thiol S-succination reaction kinetics. (b) Comparison of pH-dependent pseudo-first order rate constants for S-succination electrophiles.

The electrophilicity of maleate was less impacted by acidity (pH 5-6) than 2-bromosuccinate, consistent with the ability of protonation to increase its intrinsic reactivity (Table 1). To ensure these trends were not an artifact of our kinetic analysis method, we performed equivalent pH-dependent labeling studies in whole proteomes and made identical observations for both fumarate and maleate (Fig. S2). These studies define the chemical reactivity of the oncometabolite isomer maleate, and quantitatively validate its utility as a rapid inducer of thiol S-succination.

Inactivation of the TCA cycle by FH mutation causes a multitude of cellular effects only a subset of which may be related to protein S-succination. Previous studies have attempted to chemically induce this PTM to help isolate its influence on signaling, but have been challenged by fumarate’s low reactivity and cell permeability.^16-17^ We wondered whether the hyperreactivity of 2-bromosuccinate and maleate may overcome some of these challenges. To test this, we compared endogenous protein S-succination in FH-deficient cells (*FH -/-)* to that induced by treating an isogenic rescue (*FH* +*/*+) with 2-bromosuccinate, maleate, dimethyl fumarate, and ethyl fumarate (Fig. 4, Fig. S3). Analysis of treated cells revealed all four compounds were able to induce the PTM, as assessed by anti-S-succination immunoblotting (Fig. 4c-f). However, notable differences were also observed. Dimethyl and ethyl fumarate led to only low levels of S-succination (Fig. 4e-f). This is consistent with the chemical structures of these compounds, which would be expected to lead to the physiochemically distinct methyl- or ethyl-S-succination upon cysteine reaction.^22-23^ In contrast, 2-bromosuccinate and maleate modified a wide range of proteins, with a labeling pattern more akin to the endogenous modification found in FH -/- cells (Fig. 4c-d). Interestingly, when dosed at equivalent concentrations 2-bromosuccinate induced a stronger modification profile than maleate initially (3 h), but was superseded at extended time points. This may indicate that 2-bromosuccinate is metabolized or unstable in media over prolonged time periods.

**Figure 4.**
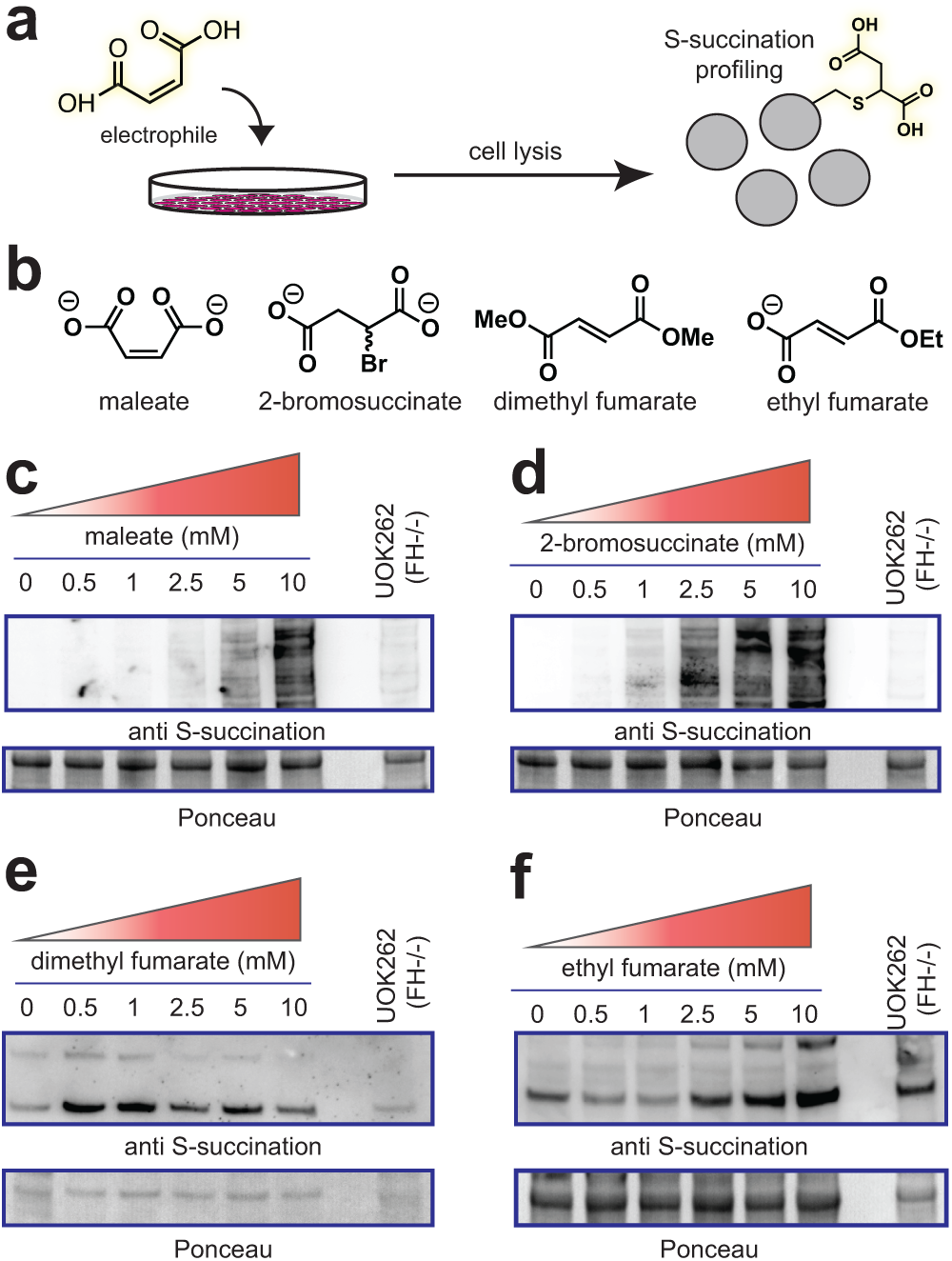
(a) Cellular evaluation of electrophilic inducers of S-succination. (b) Structure of electrophiles. (c) Cellular S-succination induced in UOK262 FH rescue (*FH*+/+) cells by maleate, (d) 2-bromosuccinate, (e) dimethyl fumarate, and (f) ethyl fumarate. Treatment time = 15 h.

During our cellular labeling studies some treatments caused noticeable detachment of cells, suggesting toxicity. As cell death may introduce a variety of biological effects that are independent of protein S-succination, we compared the cytotoxicity of fumarate, dimethyl fumarate, ethyl fumarate, methyl fumarate, 2-bromosuccinate, and maleate. HEK-293T cells were used for these studies to avoid any confounding effects FH-deficiency may have on metabolism-dependent cell death assays. We confirmed the novel reagents investigated here, 2-bromosuccinate and maleate, both cause S-succination in this cell model (Fig. S4). Analyzing dose-dependent cytotoxicity of these reagents, we found dimethyl fumarate to be the most potent inhibitor of cell growth, with a submicromolar IC50 at 48 h (Table 2, Fig. S5). This is consistent with previous studies indicating the highly electrophilic nature of this compound,^15-24^ and provides further evidence that caution should be used when interpreting biological assays using this reagent as an oncometabolite mimetic. The monoalkyl ester derivatives of fumarate, as well as 2-bromosuccinate, displayed an intermediate degree of toxicity, while fumarate and maleate were non-toxic up to millimolar concentrations. These studies specify maleate as an optimized non-toxic reagent for manipulation of cellular S-succination profiles.

As indicated above, the structure and reactivity of maleate and 2-bromosuccinate differ in subtle - but important - ways from fumarate. Therefore, as a final metric we set out to assess the effects of these two molecules on known covalent and non-covalent oncometabolite targets. As a prototypical covalent target we chose the E3 ligase KEAP1, whose covalent modification by fumarate has previously been shown to cause stabilization of the transcription factor NRF2, a master regulator of the antioxidant response.^12-13^ Consistent with the hypothesis that maleate can recapitulate aspects of fumarate’s reactivity, incubation of HEK-293T cells with maleate induced higher levels of NRF2 (Fig. 5a). To verify that this mechanism may be driven by maleate’s ability to covalently modify KEAP1, we overexpressed KEAP1 and subjected HEK-293T cells to different electrophile treatments. Transient transfection followed by lysis, KEAP1 immunoprecipitation, and anti-S-succination immunoblotting revealed detectible covalent modification of KEAP1 upon addition of maleate (Fig. 5b). 2-bromosuccinate also was able to trigger NRF2 stabilization and KEAP1 S-succination, although cytotoxicity was observed at the higher dosage (10 mM; Fig. 5c). In contrast, DMF and fumarate itself did not induce detectable S-succination of KEAP1, presumably due to a lack of ester hydrolysis and cell permeability, respectively. Turning our attention to non-covalent targets of fumarate, it has previously been shown that inactivation of FH in HLRCC causes accumulation of the transcription factor HIF-1α, which is attributed to fumarate’s ability to competitively inhibit KG-dependent prolyl hydroxylases involved in HIF-1α degradation.^6-7^ In contrast to its effects on NRF2, incubation of HEK-293T cells with maleate (1-10 mM) for 24 h did not cause a substantial increase in HIF-1α levels. However, incubation with 2-bromosuccinate caused a detectible increase in HIF-1α levels (Fig. 5a). This suggests that 2-bromosuccinate - or one of its metabolites - may function as a competitive inhibitor of prolyl hydroxylase activity, while maleate cannot. Previous crystallographic studies of prolyl hydroxylases complexed with a KG-competitive ligands are more consistent with a trans-orientation near the Fe(II)-KG reaction center, which cannot be accessed by cis-fumarate.^25-26^ Together, these data suggest the unique structure of maleate may in certain instances allow selective induction of fumarate’s signature non-enzymatic PTM (S-succination) while limiting its non-covalent effects on KG-dependent enzymes such as prolyl hydroxylases.

**Figure 5.**
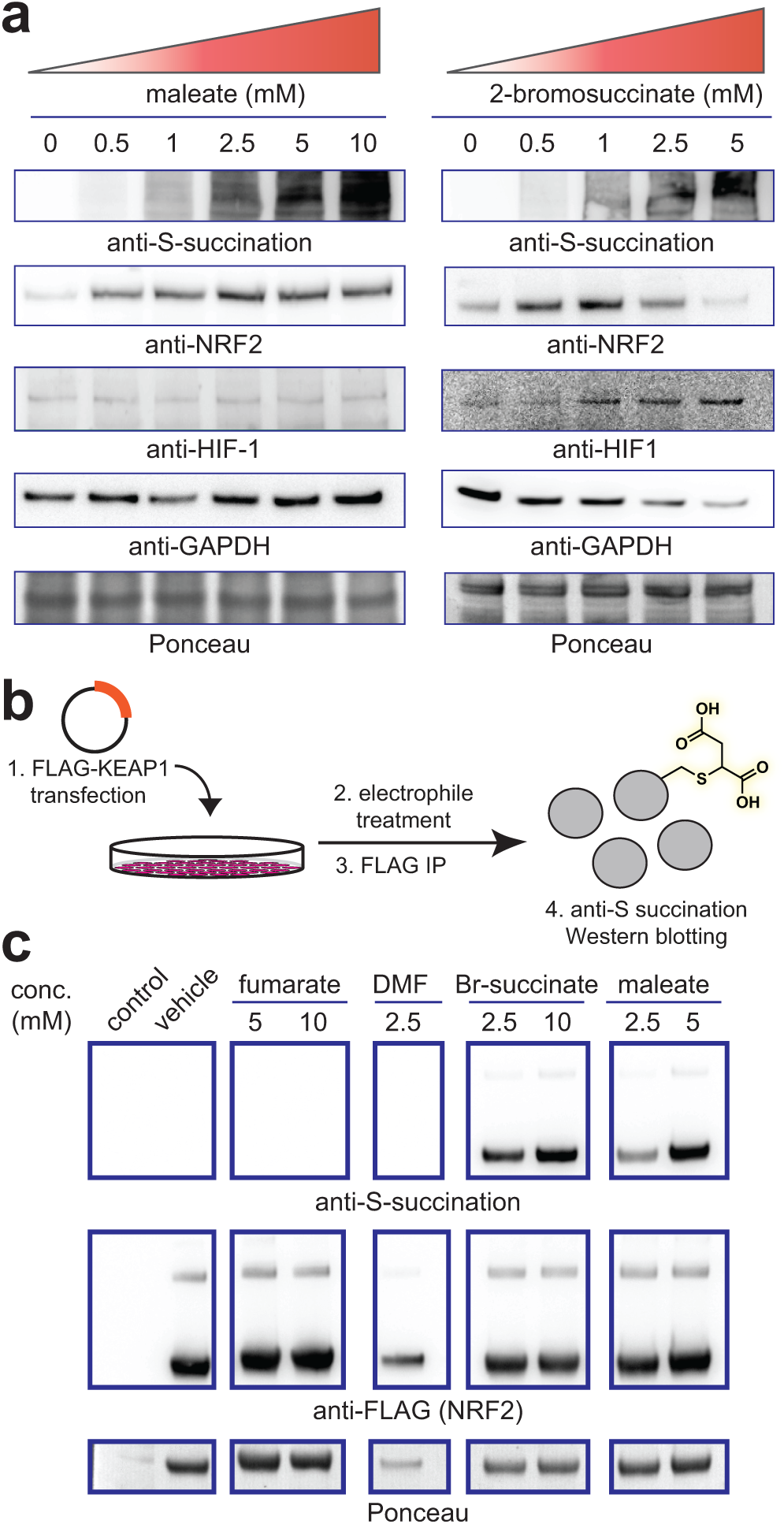
(a) Dose-dependent effects of maleate and 2-bromosuccinate on S-succination, NRF2, and HIF1-a in HEK-293T cells. Treatment time = 15 h. (b) Assay for assessing cellular S-succination of ectopically-expressed FLAG-KEAP1. (c) Cellular S-succination of ectopically-expressed KEAP1 by electrophiles (treatment time = 15 h). Br-succinate = 2-bromosuccinate. DMF = dimethyl fumarate.

Recent advances in chemoproteomics have enabled the detection and comprehensive profiling of metabolite-protein interactions.^27-31^ Translating these datasets into mechanistic knowledge requires methods to manipulate these interactions in living cells, but is challenged by the limited permeability and tempered reactivity of many metabolites. Here we report a new approach for the induction of cysteine S-succination, a non-enzymatic PTM produced by the covalent oncometabolite fumarate. In contrast to previous methods that utilize fumarate esters, the approach reported here recapitulates the endogenous PTM (rather than an ester-linked version) and can be applied in living cells with limited toxicity. Furthermore, our initial studies indicate that in HEK-293T cells one of these reagents, maleate, is able to differentially induce covalent (KEAP1 S-succination/NRF2 activation) versus non-covalent (PHD inhibition/HIF-1α stabilization) effects of fumarate. This property should be useful in studies seeking to define the relative influence of these two mechanisms on the plethora of biological phenomena in which fumarate has been implicated in HLRCC, which include altered gene expression, mitotic entry, and epithelial to mesenchymal transition.^5-32-33^ Finally, we note some limitations of our method as currently comprised. First, while maleate, 2-bromosuccinate, and fumarate induce the same PTM, they differ substantially in their reactivity and most notably show distinct pH-dependent labeling profiles. Our previous studies have provided evidence that hydrogen or dihydrogen fumarate is the reactive species underlying covalent S-succination in HLRCC,^14^ and if the S-succination agonists reported here are reactive in the absence of protonation this may alter their proteomic reactivity profile. Future comparisons of these reagents and fumarate in competitive chemoproteomic experiments will help define the similarity (or distinctiveness) of their cysteine reactivity landscapes. Another key difference between FH-deficiency and this exogenous S-succination strategy is that in the former fumarate diffuses out of the mitochondria, while in the latter electrophiles diffuse in through the plasma membrane. Targeted delivery of maleate or 2-bromosuccinate to the mitochondria may be required to enable the study of S-succination in this organelle.^34-35^ In the immediate future, we anticipate the less toxic maleate reagent will be useful for determining whether S-succination elicits a distinct gene expression response relative to other NRF2-inducing electrophiles,^5^ as well as in isotopic labeling strategies designed to assess S-succination stoichiometry.^36^ In the longer term, we envision maleate’s unique status as an anion electrophile that is relatively insensitive to bulk pH may prove useful in the construction of covalent fragment libraries designed to interrogate the ligandability of distinct subsets of the proteome.^37^ By expanding our inventory of methods for the study of metabolite-derived PTMs, these studies provide a foundation for defining and manipulating the signaling role of metabolism in cancer.

## ASSOCIATED CONTENT

Figures S1-S5 and supporting materials and methods. This material is available free of charge via the Internet.

## ACKNOWLEDGMENT

This work was supported by the Intramural Research Program of NIH, the National Cancer Institute, The Center for Cancer Research (ZIA BC011488-04).

