## Supplemental Information for "An Oncometabolite Isomer Rapidly Induces A Pathophysiological Protein Modification"

**Table of Contents for Supporting Information**

| | Page |
|---|---|
| Table of Contents | S1 |
| Supplementary Figures S1-S5 | S2-6 |
| General materials and methods | S7 |
| Cell culture and isolation of whole-cell lysates | S8 |
| In vitro S-succination assays | S9 |
| Kinetic analysis of thiol S-succination | S10 |
| Cellular S-succination assays | S11 |
| Cytotoxicity assays | S12 |
| Analysis of KEAP1 S-succination | S13 |
| Full gels and blots | S14-29 |
| References | S30 |

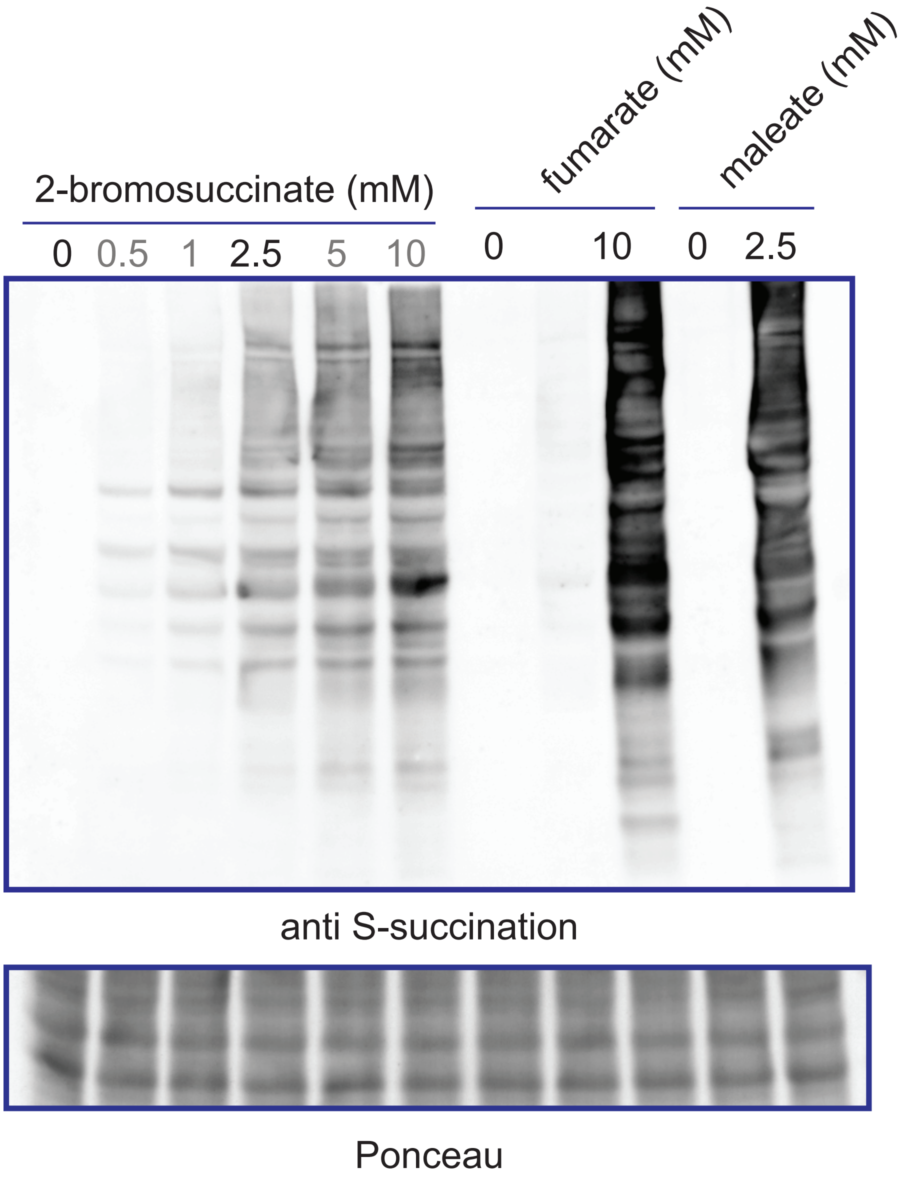


**Figure S1** Direct comparison of relative in vitro labeling of HEK-293T cell lysates by 2-bromosuccinate, fumarate, and maleate. Proteomes were incubated with each reagent at the specified concentration for 15 hours.

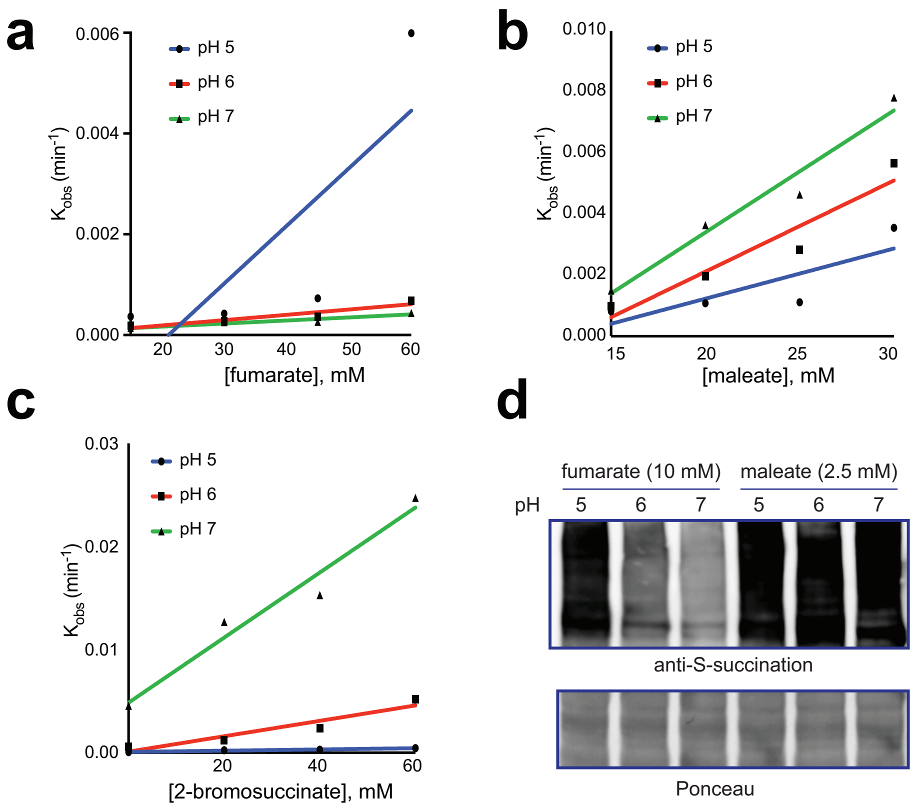


**Figure S2** Kinetic analysis of 2-thiophenol S-succination by (a) fumarate, (b) maleate, and (c) 2-bromosuccinate. (d) Analysis of pH-dependent S-succination of HEK-293T cell protemes by fumarate and maleate. Proteomes were incubated with each reagent at the specified concentration for 15 hours.

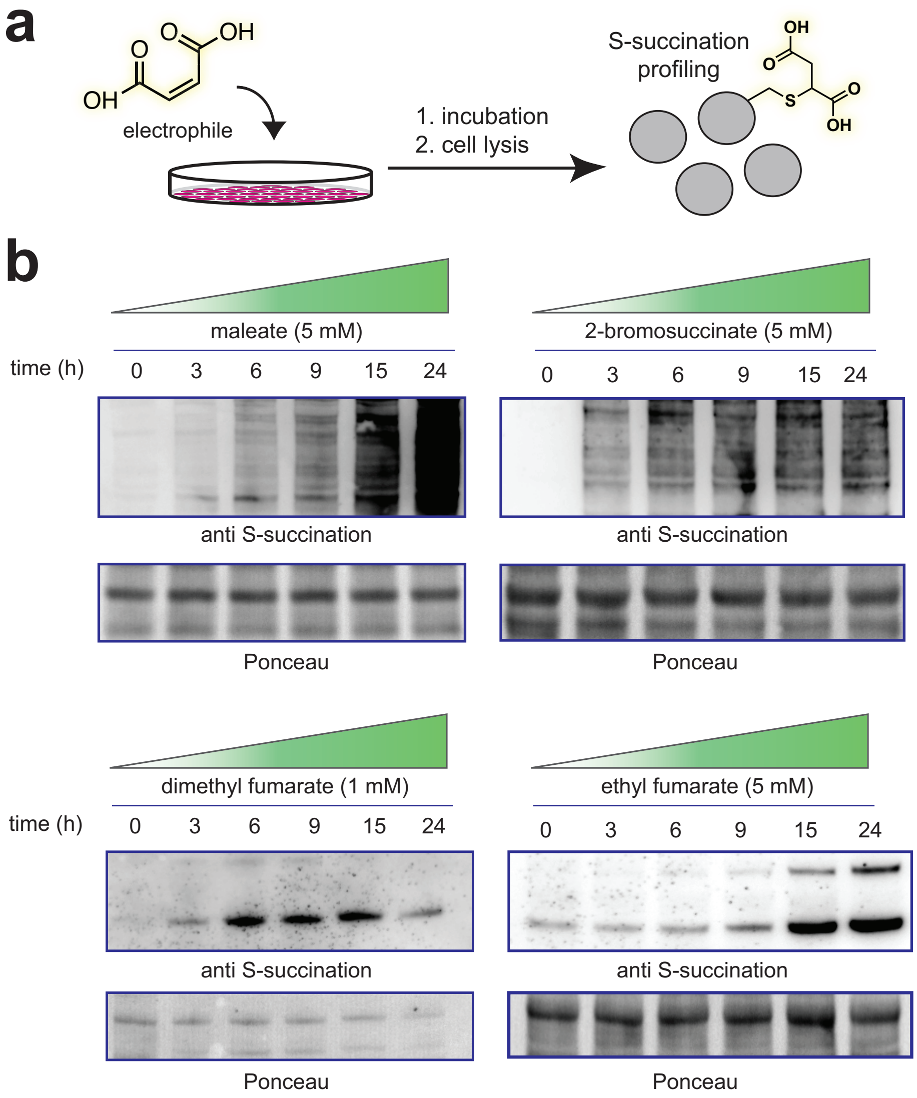


**Figure S3** (a) Schematic for chemical induction of S-succination in living cells. (b) Time-dependent S-succination profiles of maleate, 2-bromosuccinate, dimethyl fumarate, and ethyl fumarate in UOK262 *FH+/+* ("FH rescue") cell lines.

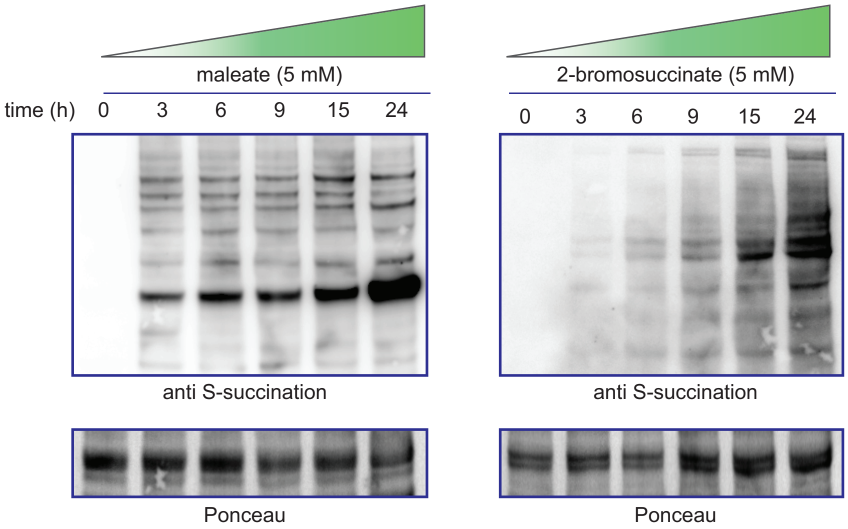


**Figure S4** Treatment of HEK-293T cells with maleate (left) or 2-bromosuccinate leads to time-dependent proteomic S-succination.

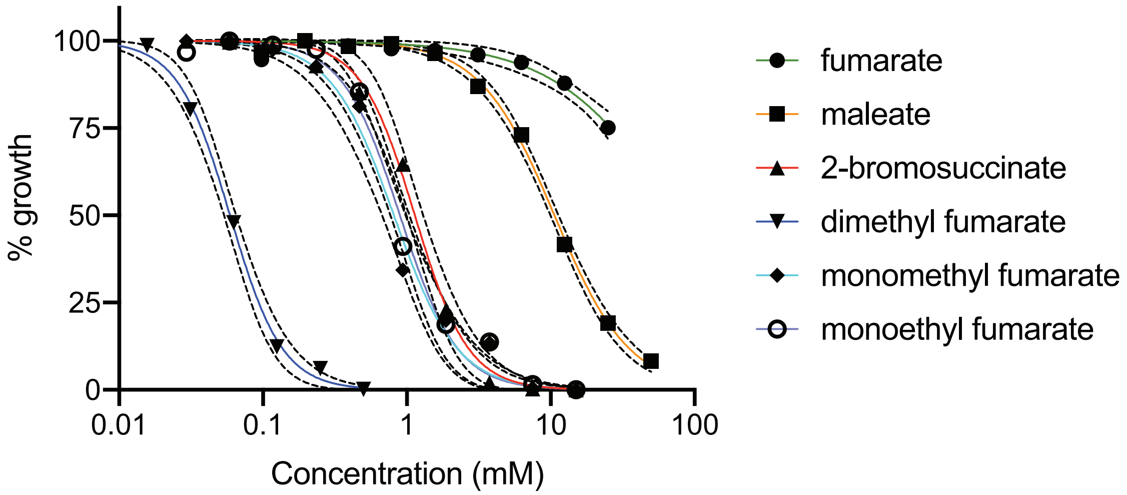


**Figure S5.** Relative cytotoxicity of cellular S-succination reagents in HEK-293T cells. Cells were treated with compounds for 48 h.

**General materials and methods**

Unless otherwise specified all chemicals were used without further purification. Fumaric acid (A10976) and ethyl fumarate (A12545) were purchased from Alfa Aesar. Maleic acid (M0375), bromosuccinic acid (B81204), and dimethyl fumarate (242926) were purchased from Sigma. For western blotting, SDS-PAGE was performed using Bis-Tris NuPAGE gels (4-12%, Invitrogen #NP0322), and MES running buffer (Life technologies #NP0002) in Xcell SureLock MiniCells (Invitrogen) according to the manufacturer's instructions. Gels were transferred to nitrocellulose membranes (Novex, Life Technologies # LC2001) by electroblotting at 30 volts for 1 hour using a XCell II Blot Module (Novex). Membranes were blocked using StartingBlock (PBS) Blocking Buffer (Thermo Scientific) for 30 minutes, then incubated overnight at 4°C in a solution containing the primary antibody of interest (1:200 dilution for S-(2-succinyl)-cysteine antibody and 1:1000 dilution for all other antibodies) in the above blocking buffer with 0.05% Tween 20. The membranes were next washed with TBST buffer, and incubated with a secondary HRP-conjugated antibody (anti-rabbit IgG, HRP-linked [7074], Cell Signaling, 1:1000 dilution) for 1 hour at room temperature. The membranes were again washed with TBST, treated with chemiluminescence reagents (Western Blot Detection System, Cell Signaling) for 1 minute, and imaged for chemiluminescent signal using an ImageQuant Las4010 Digitial Imaging System (GE Healthcare). All blots were imaged using 20X LumiGLO® Reagent and 20X Peroxide (Cell Signaling Technologies #7003) except the anti-S-(2-succinyl)-cysteine antibody which used SuperSignal (Thermo Fisher #37074). Cysteine S-succination antibody was purchased from Discovery Antibodies (crb2005017).[1] FLAG (2044S), HIF1-a (3716S), and GAPDH (5174S) antibodies were purchased from Cell Signaling. NRF2 antibody was purchased from Novus (NBP1-32822). Flag-Keap1 plasmid was a gift from Qing Zhong (Addgene plasmid # 28023). Anti-FLAG pulldown was performed using immunoprecipitation kit (Sigma - FLAGIPT1) from Sigma. HEK-293T cells were obtained from the NCI Tumor Cell Repository. UOK262 (*FH-/-*), UOK262WT (*FH +/+* rescue) cells were obtained from Linehan lab.[2] Transfection quality plasmid was generated using the PureLink™ HiPure Plasmid Maxiprep Kit (Thermo Fisher # K210006). Qubit Protein Assay kit was purchased from Thermo Fisher #Q33211.HPLC analyses of electrophile reactivity were performed using an Agilent 1250 Infinity HPLC equipped with a UV detector and a Kinetex C18 column (100 x 2.1 mm, 100 Å, 2.6 μm). Data analysis was performed using GraphPad Prism v7.0 software.

**Cell culture and isolation of whole-cell lysates**

All cell lines were cultured at 37 °C under 5% CO 2 atmosphere in a growth medium of RPMI supplemented with 10% FBS and 2 mM glutamine, with the exception of UOK262 cell lines, which were cultured in DMEM supplemented with 10% FBS, 2 mM glutamine, 1 mM pyruvate and 0.3 mg/mL of G418. Cells were dosed with electrophiles by adding stock solutions (1000× stock in DMSO or water) directly to growth medium at the specified concentration. Unfractionated proteomes were harvested from cell lines (80–90% confluency) by scraping cells into a Falcon tube and centrifuging (500 g × 3 min, 4 °C) to form a cell pellet. After removal of PBS supernatant, cell pellets were washed ice cold PBS, and either stored at −80 °C or immediately lysed by sonication. For lysis, cells were first resuspended in 40 uL ice cold PBS (10–20 × $10^6$ cells/mL) containing protease inhibitor cocktail (1X, EDTA-free, Cell Signaling Technology # 5871S). These samples were then lysed by sonication using a 100 W QSonica XL2000 sonicator (3 × 1s pulse, amplitude 1, 60s resting on ice between pulses). Lysates were pelleted by centrifugation (14,000 rcf × 30 min, 4 °C) and quantified on a Qubit 2.0 Fluorometer using a Qubit Protein Assay Kit. Quantified proteomes were diluted to 2 mg/mL and analyzed immediately or stored at −80 °C for analysis.

**In vitro S-succination assays**

Dose-dependent induction of S-succination was tested by treating isolated proteomes (40 μg) with maleate and 2-bromosuccinate for 15 hours. Time-dependent induction of S-succination was tested by treating isolated proteomes (40 μg) with maleate and 2-bromosuccinate for for 0, 3, 6, 9, 15, or 24 hours at the specified concentration. Reaction pH was controlled via neutralization with NaOH or HCl. Reactions were quenched with 4x SDS-PAGE loading buffer containing 10% DTT, and care was taken not to boil samples prior to gel loading so as not to induce artifactual S-succination. For all in vitro/proteomic S-succination analyses, samples were loaded onto a SDS-PAGE gel and analyzed by Western blot as described above using S-(2-succinyl)-cysteine antibody (Discovery Antibodies) at 1:200 dilution factor and imaged using SuperSignal (Thermo Fisher).

**Kinetic analysis of thiol S-succination**

*HPLC Analysis*

Kinetics of thiophenol succination were analyzed as previously described.[3] Briefly, 10 uL 60 mM thiophenol (Sigma Aldrich) in ACN, 10 uL electrophile (fumaric, maleic, or bromosuccinic acid) in DMSO, 2 uL 10 mM 7-diethylamino-4-methylcoumarin (Alfa Aesar) in DMSO, and 5 uL 400 mM tris(hydroxymethyl)phosphine (Acros Organics) in water were combined and adjusted to a final volume of 200 uL with PBS pH 7.2 (Gibco). Solution pH was adjusted to 5, 6, and 7 by adding 2 M NaOH or 1 M HCl dropwise as necessary and checked with pH strips (Millipore MColorpHast pH 2.0-9.0). 10 uL reaction mixture was injected into an Agilent Technologies 1260 Infinity HPLC equipped with a UV detector and separated using a Kinetex 2.6 um C18 100A 100x2.1 mm column at a flow rate of 0.5 mL/min. UV detector was set at 254 nm. Buffer A: 0.1% TFA, Buffer B: ACN with a gradient as follows:

| Time (min) | % A | % B |
|---|---|---|
| 0 | 100 | 0 |
| 2 | 100 | 0 |
| 18 | 50 | 50 |
| 20 | 0 | 100 |
| 23 | 0 | 100 |
| 26 | 100 | 0 |
| 30 | 100 | 0 |

*Kinetic Analysis*

Appearance of the product peak (as determined by injecting a succinated standard prior to kinetic analysis) was monitored over a 15 hour period relative to the internal standard. Kinetics were calculated according to the method of Gupta et al.[4] In brief, area under the product peak was normalized as percent of area under the internal standard peak and data were fit to a one phase exponential decay equation. The data were constrained to a plateau of the observed highest value for each electrophile and the equation used was $Y = (Y_0 - 8.00)*e^{(-k*t)} + B$; where Y is the normalized absorbance at a given time, $Y_0$ is the initial absorbance, t is time in minutes, and k is the apparent rate constant for the succination reaction at the given pH (in units of $hr^{-1}$), and B is the plateau percentage value. k for each reaction condition was plotted against pH and a linear regression was performed, the slope of which was considered the pseudo-first order rate constant for the reaction.

### Cellular analysis of cysteine S-succination

HEK293T and UOK262 *FH+/+* ("FH rescue") cells were plated in 6-well dishes at 7 $\times 10^5$ and 3 $\times 10^5$ cells per well, respectively. Cells were allowed to adhere for 24 hours before dosing with electrophilic S-succination reagents. Dose-dependent induction of S-succination was tested by incubating cells with the specified concentration of each electrophile (0-10 mM) for 15 hours. Time-dependent induction of S-succination was tested by incubating cells with the specified concentration of each electrophile for 0, 3, 6, 9, 15, or 24 hours. After incubation, cells were harvested, and soluble proteome was isolated and quantified as described above. For western blot analysis, 20 µg of lysates were loaded per lane of the gel. For all cellular S-succination analyses, samples were loaded onto a SDS-PAGE gel and analyzed by Western blot as described above using S-(2-succinyl)-cysteine antibody (Discovery Antibodies) at 1:200 dilution factor and imaged using SuperSignal (Thermo Fisher).

### Cytotoxicity assays

Cytotoxicity of electrophilic S-succination inducting reagents was measured by assessing cellular ATP content using Celltiter Glo 2.0 (Promega, 7570). Briefly, stock solutions of maleate, fumarate, and 2-bromosuccinate stock solutions were prepared in equimolar base solutions and pH neutralized. Ethyl fumarate, monomethyl fumarate, and dimethyl fumarate stock solutions were prepared in neat DMSO to avoid hydrolysis, and dosed to give a final concentration of ≤ 0.1% DMSO. HEK-293T cells were plated in 96-well plates in 50 µL of the defined media at 8 $\times 10^3$ cells/well. After 24 hours, electrophiles were added to adhered cells in 50 µL of media and serially diluted. Quadruplicate wells were used for each stock concentration. After 48 hours, Celltiter Glo Reagent 2.0 was prepared and added to the well in a 1:1 ratio (100 uL of cells + 100ul of Celltiter Glo), giving a final well volume of 200 uL per well. Plates were placed in a shaker and incubated at room temperature for 30 minutes before measurement of luminescence was on an Enspire microplate reader (PerkinElmer). Data are charted as a percentage of cell viability compared to untreated controls, corrected for background luminescence. IC50 was defined as the concentration that inhibits 50% of control cell growth and determined by non-linear least-squares regression fit to $Y = A + (B - A)/(1+10 \curlywedge ((\text{Log EC50} - X) \times H$, where $A$ = max., $B$ = min. and $H$ = Hill Slope. All calculations were performed using Prism 4 (GraphPad) software. IC50 values represent the mean and standard deviation of three independent biological replicates.

**Analysis of KEAP1 S-succination**

HEK-293T cells were plated in 10 cm dishes (3 x$10^6$ cells/dish) and allowed to adhere and grow for 24 hours. Cells were transfected with 20 μg KEAP1-FLAG using Lipofectamine 2000 (Invitrogen # 11668019) and allowed to incubate for 48 hours for overexpression analyses. To assess the ability of electrophiles to induce S-succination of KEAP1, electrophiles were added to transfected cells exactly 33 hours following transfection and allowed to incubate for an additional 15 hours. For both vehicle and treated samples, cells were harvested 48 hours after transfection, and lysed in ice-cold PBS and protease inhibitor cocktail as descrbied above. Cell lysates were quantified, diluted to 1.5mg/mL and incubated with the anti-FLAG affinity resin over night at 4°C. Washes and anti-FLAG elutions were carried out according to manufacturer's instructions. Eluted protein was analyzed by SDS-PAGE and subjected to immunoblotting and probing using anti-FLAG and anti-2-succynyl-cysteine antibodies as described above.

**Full gels and blots**

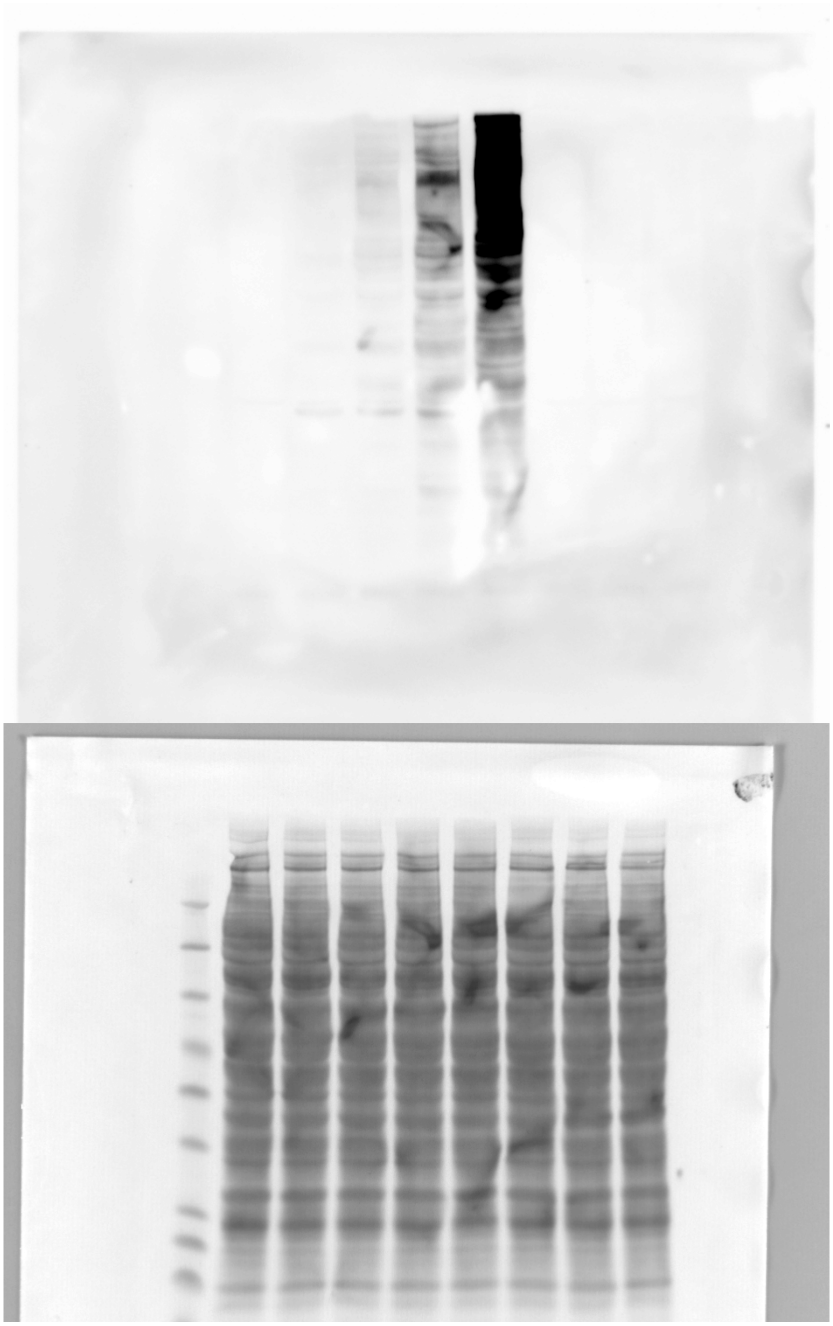

Full blot for Figure 2b, dose-dependent HEK-293T proteome maleate treatments (lanes 1-5).
Top: Chemiluminescence. Bottom: Ponceau.

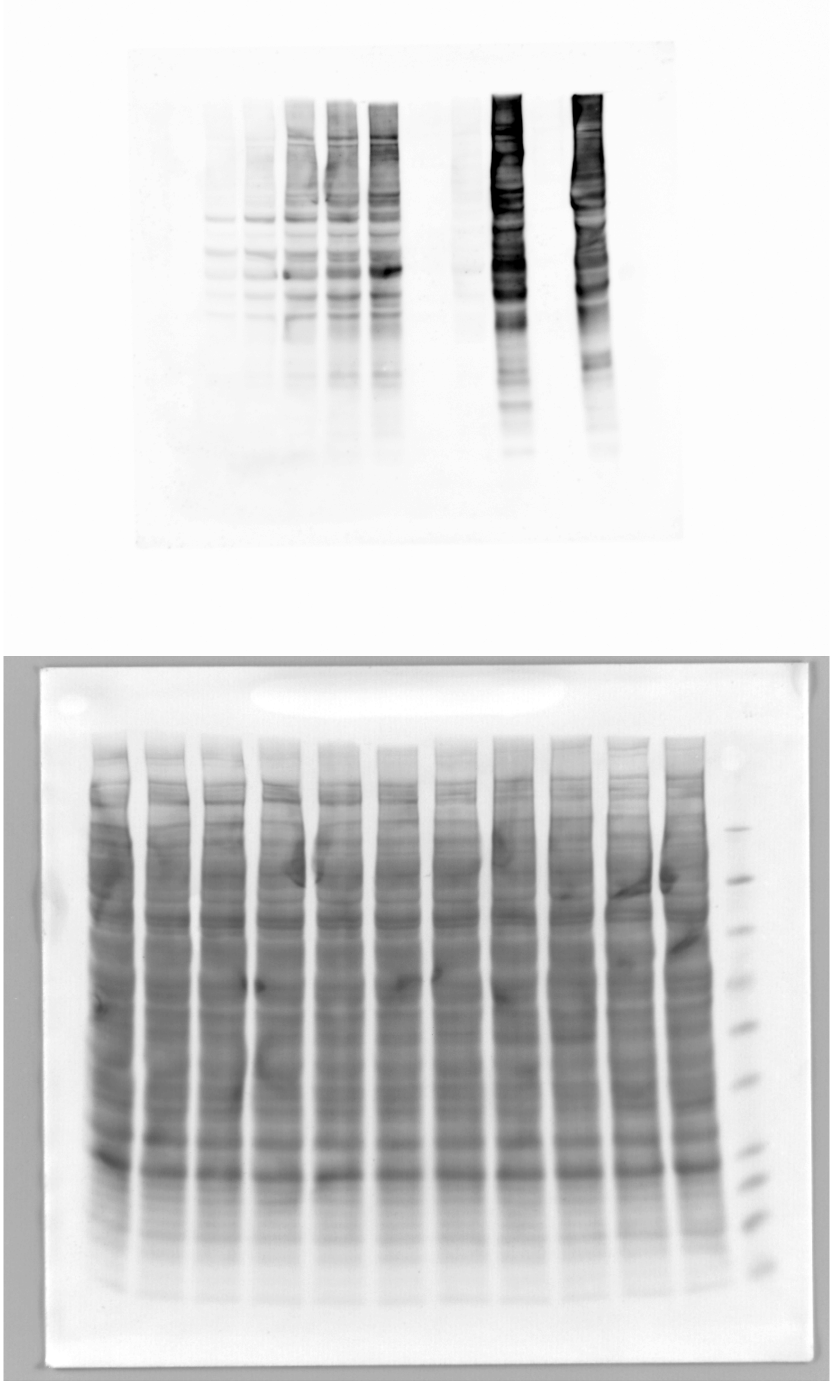

Full blot for Figure 2b and Figure S1, dose-dependent 2-bromosuccinate (lanes 1-6), fumarate (lanes 7-9), and maleate (lanes 10-11) treatments. Top: Chemiluminescence. Bottom: Ponceau.

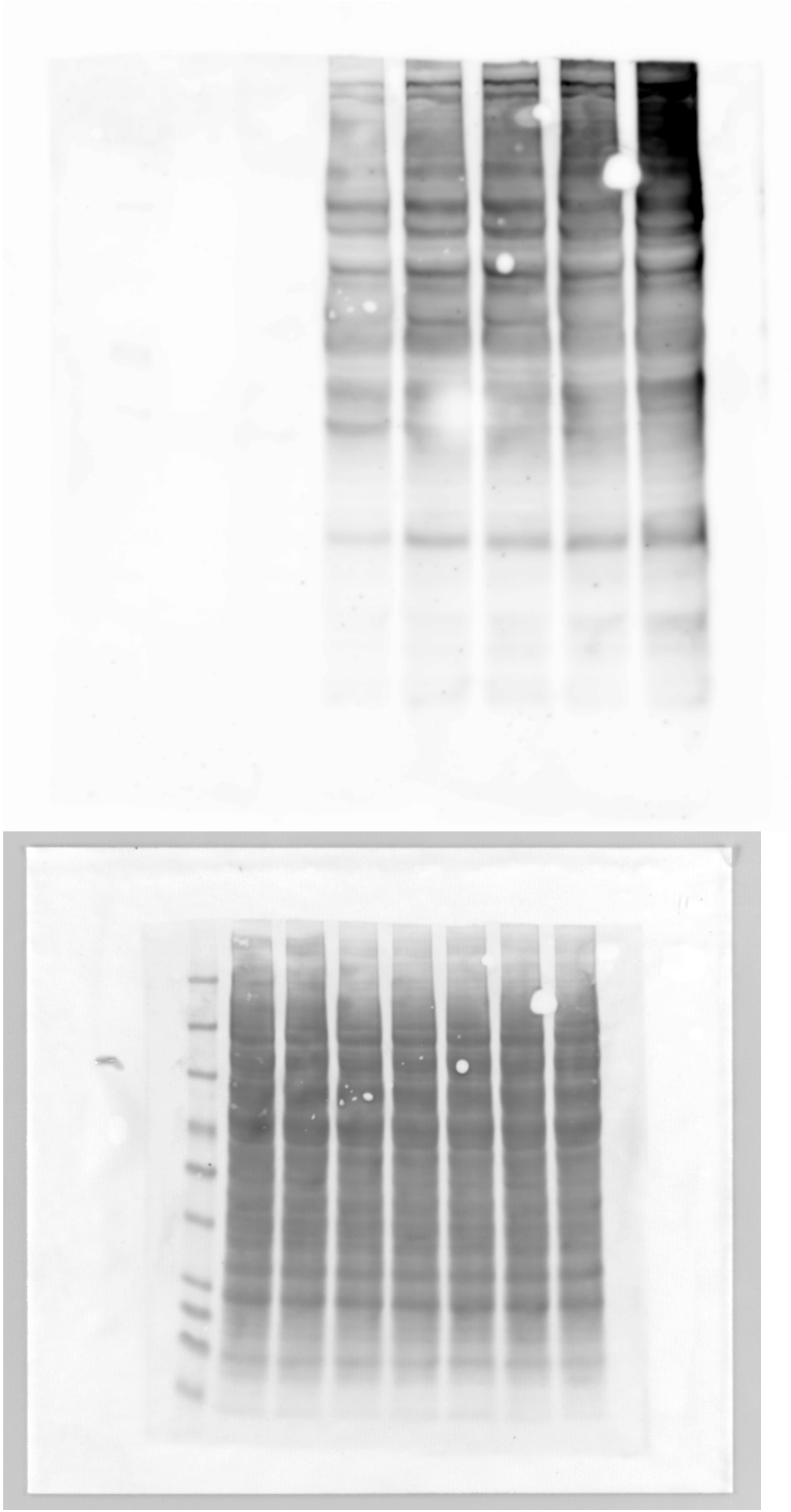

Full blot for Figure 2c, time-dependent HEK-293T proteome maleate treatments (lanes 1-7).
Top: Chemiluminescence. Bottom: Ponceau.

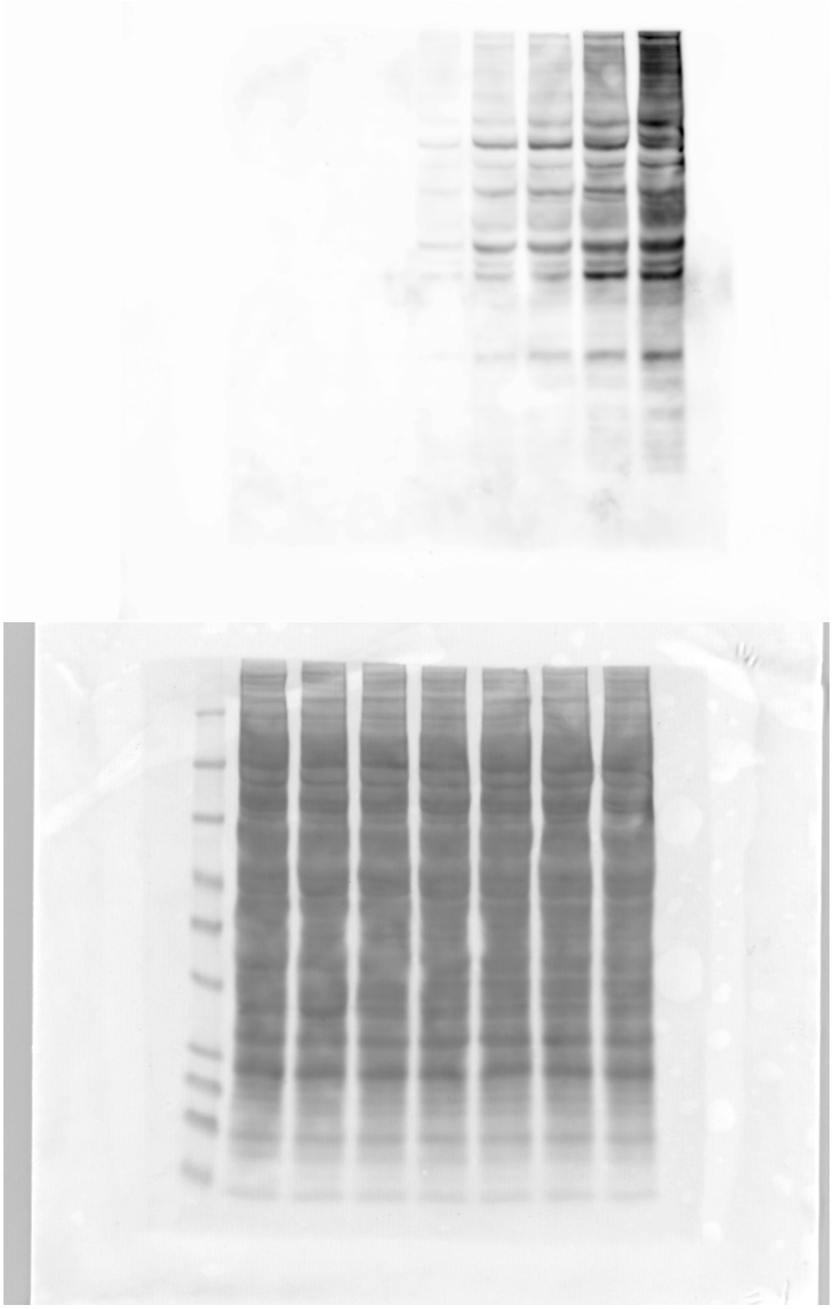

Full blot for Figure 2c, time-dependent HEK-293T proteome 2-bromosuccinate treatments (lanes 1-7). Top: Chemiluminescence. Bottom: Ponceau.

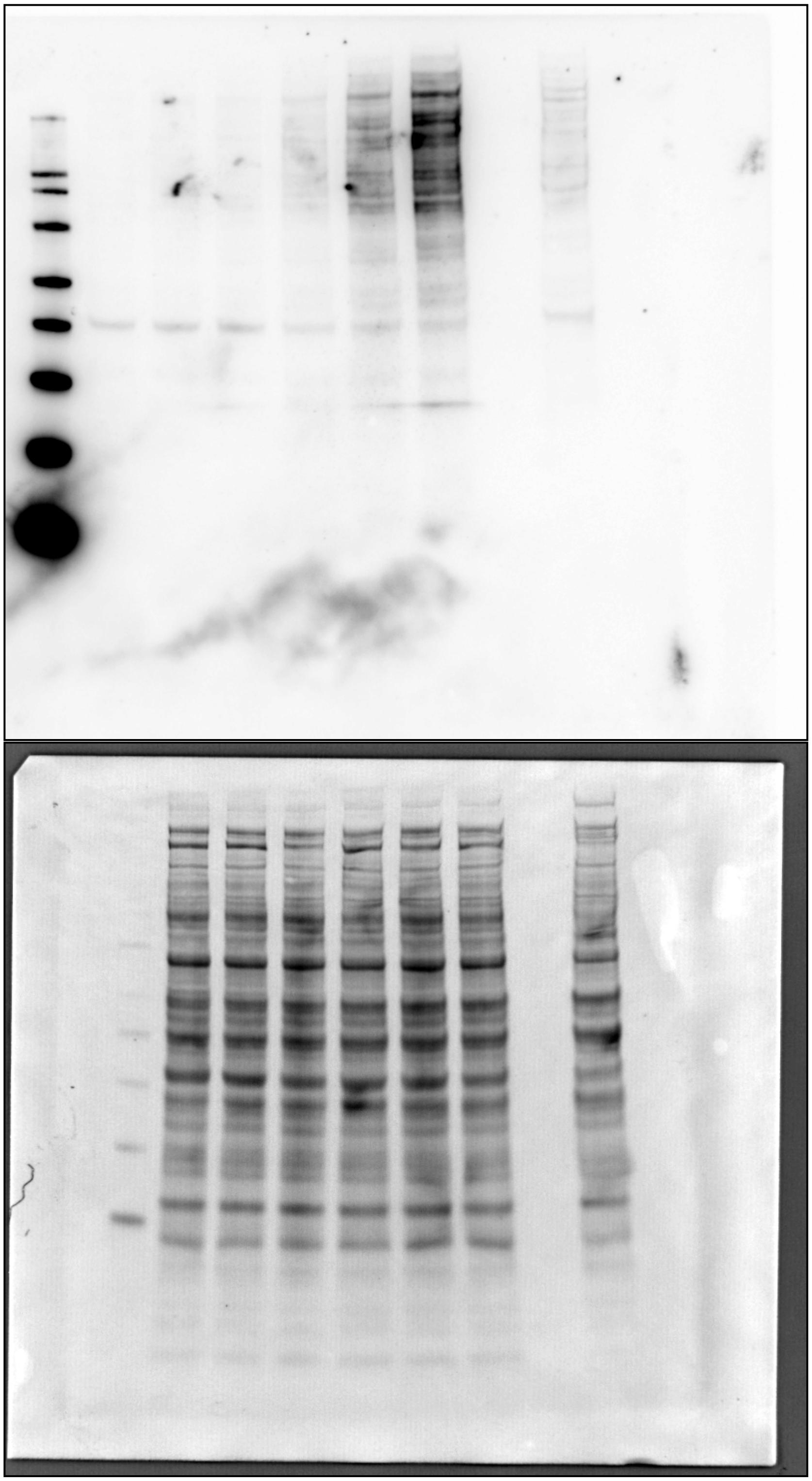

Full blot for Figure 4c, dose-dependent cellular S-succination by maleate (lanes 2-7). Top: Chemiluminescence. Bottom: Ponceau.

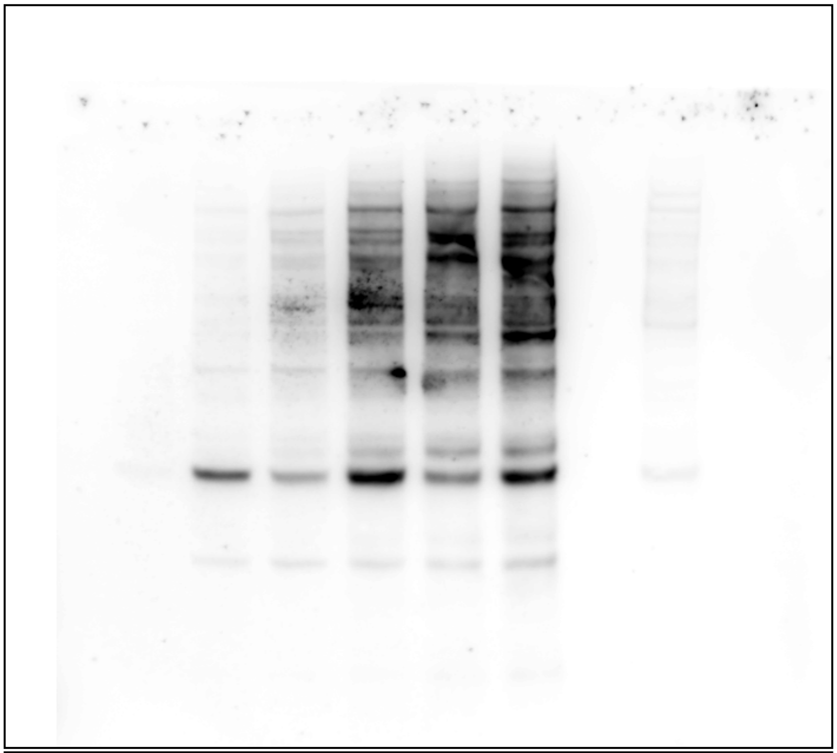

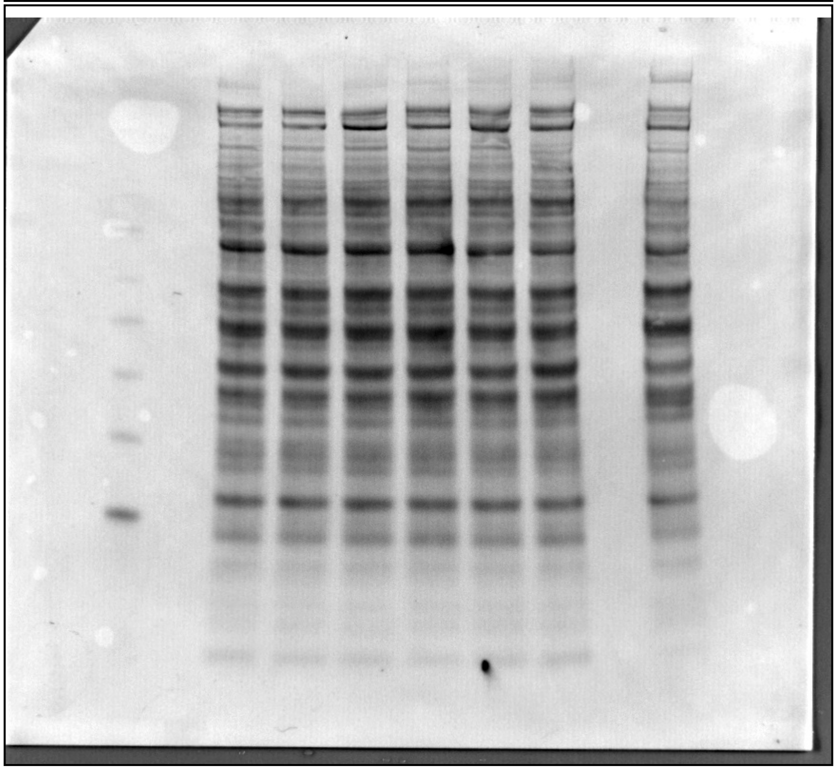

Full blot for Figure 4d, dose-dependent cellular S-succination by 2-bromosuccinate (lanes 1-6 top, lanes 3-8 bottom). Top: Chemiluminescence. Bottom: Ponceau.

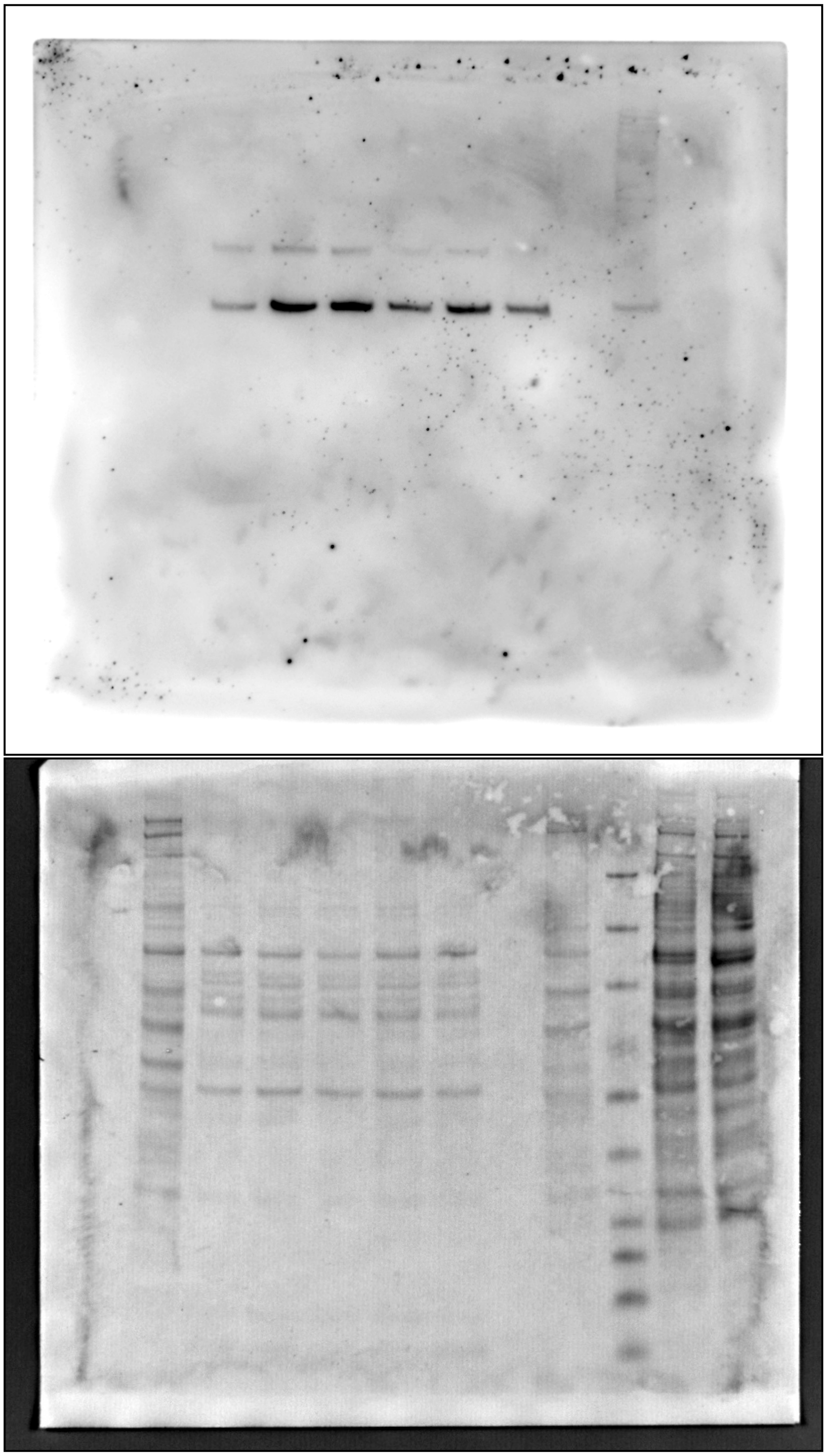

Full blot for Figure 4e. Dose-dependent cellular S-succination by dimethyl fumarate. Top: Chemiluminescence. Bottom: Ponceau.

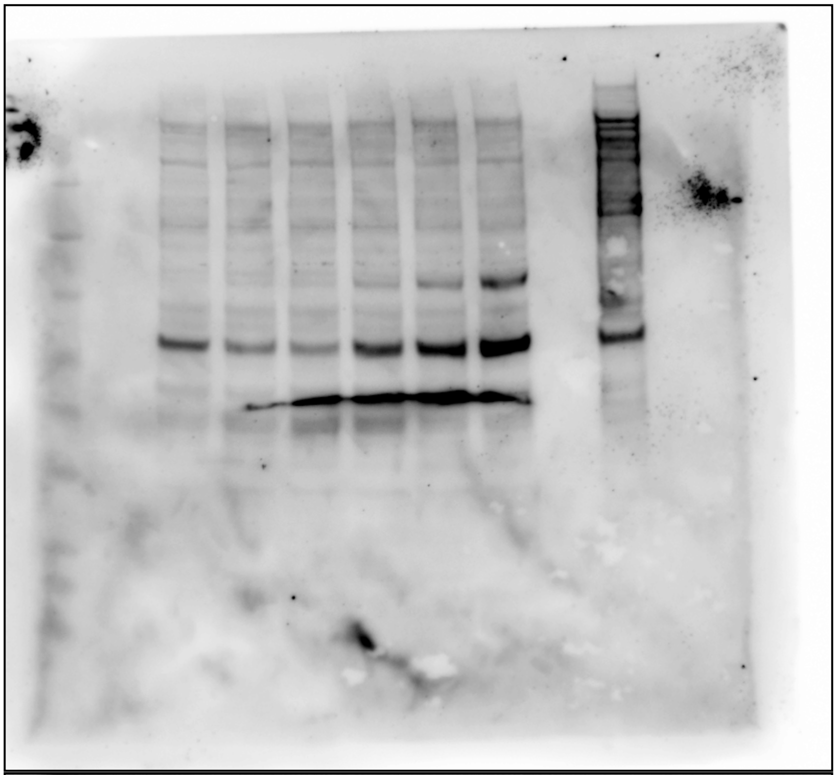

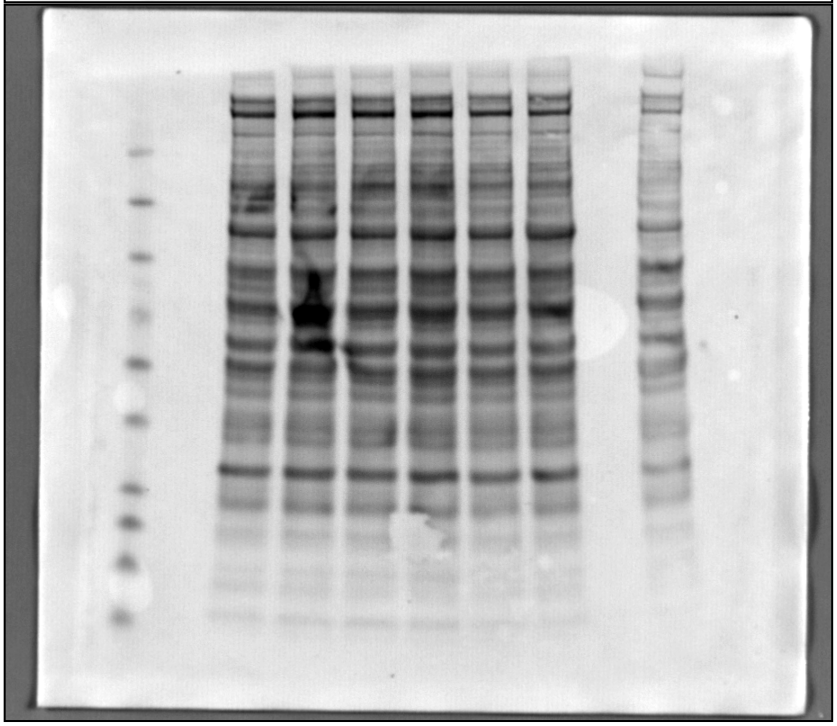

Full blot for Figure 4f. Dose-dependent cellular S-succination by ethyl fumarate. Top: Chemiluminescence. Bottom: Ponceau.

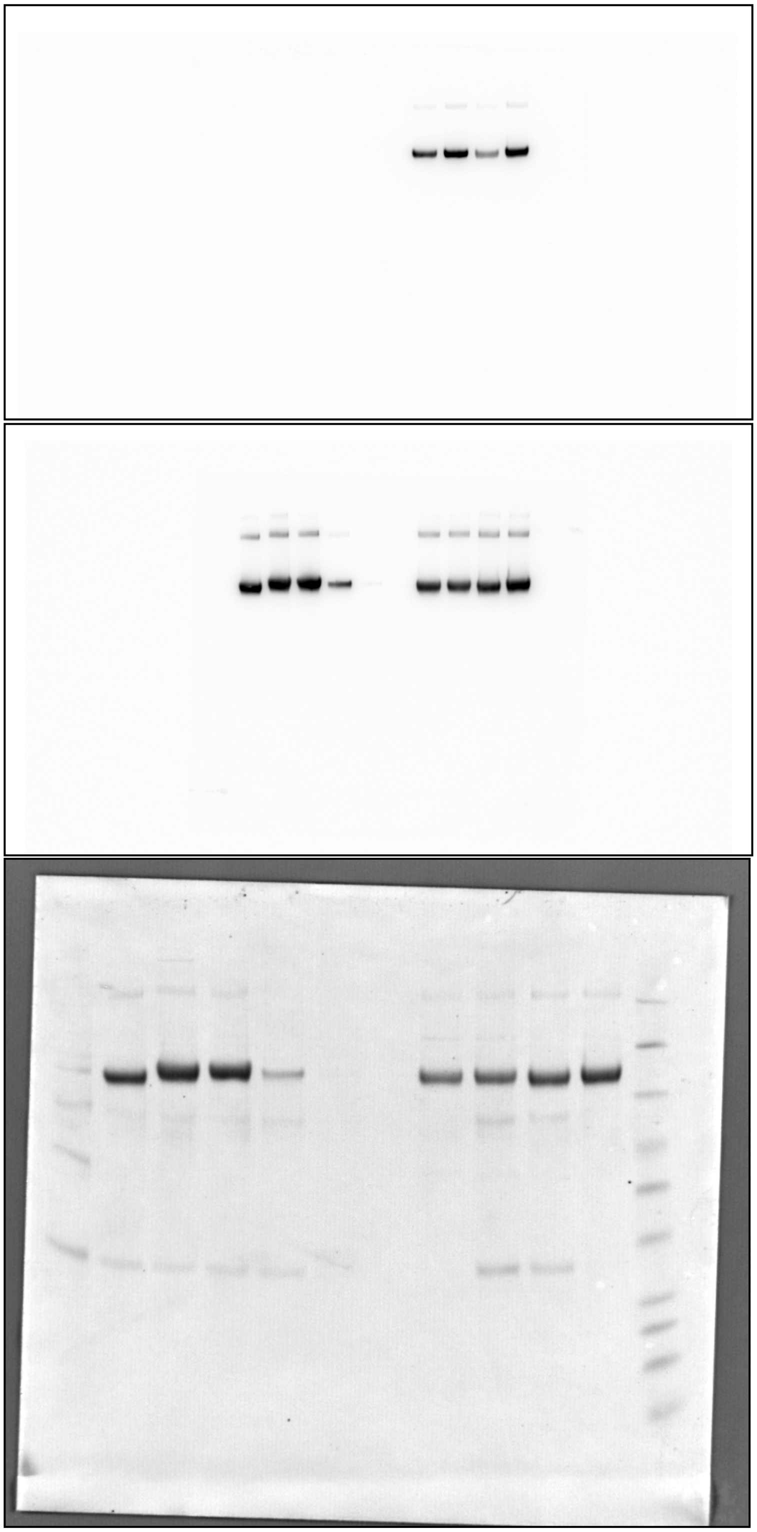

Full blot for Figure 5c. Top: Chemiluminescence, anti-S-succination. Middle: Chemiluminescence, anti-FLAG. Bottom: Ponceau.

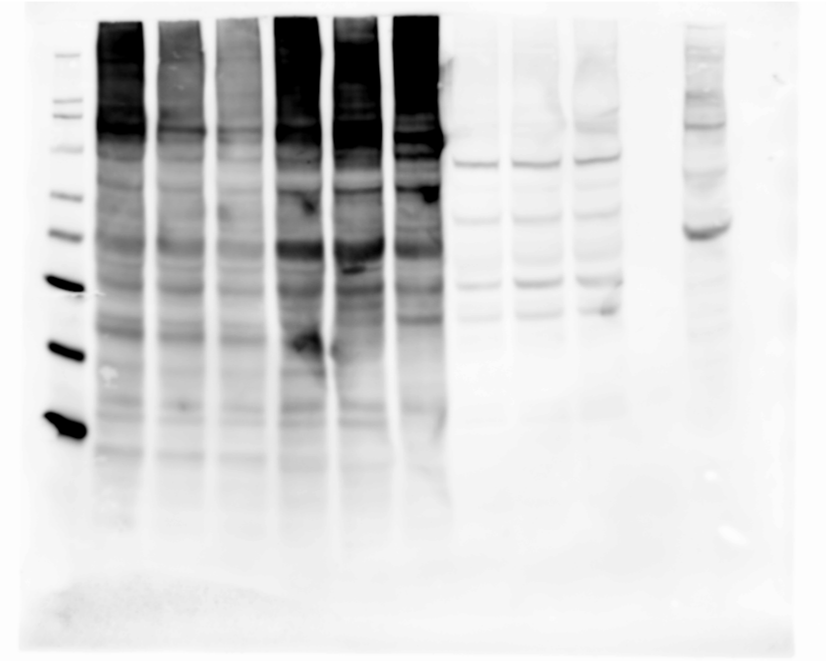

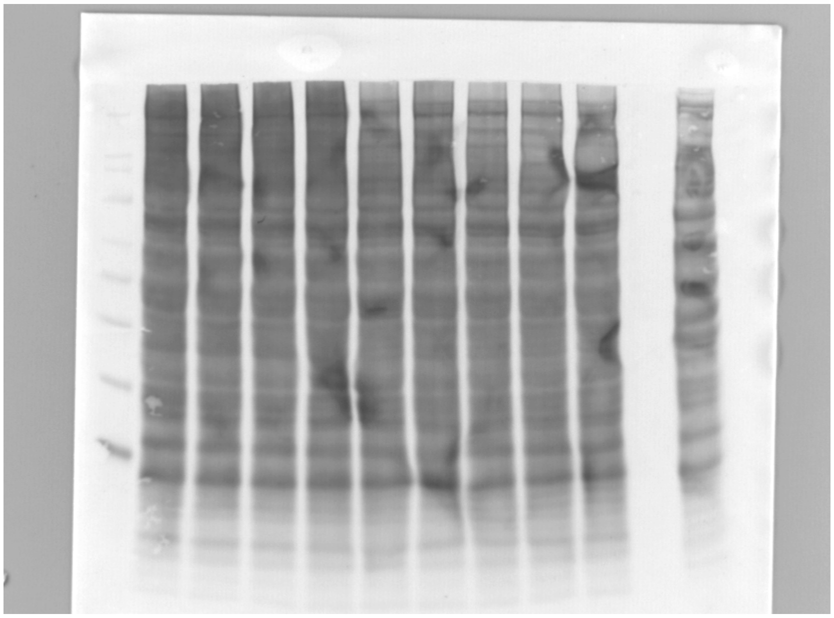

Full blot for Figure S2d, pH-dependent S-succination by fumarate (lanes 2-4) and maleate (lanes 5-7). Top: Chemiluminescence. Bottom: Ponceau.

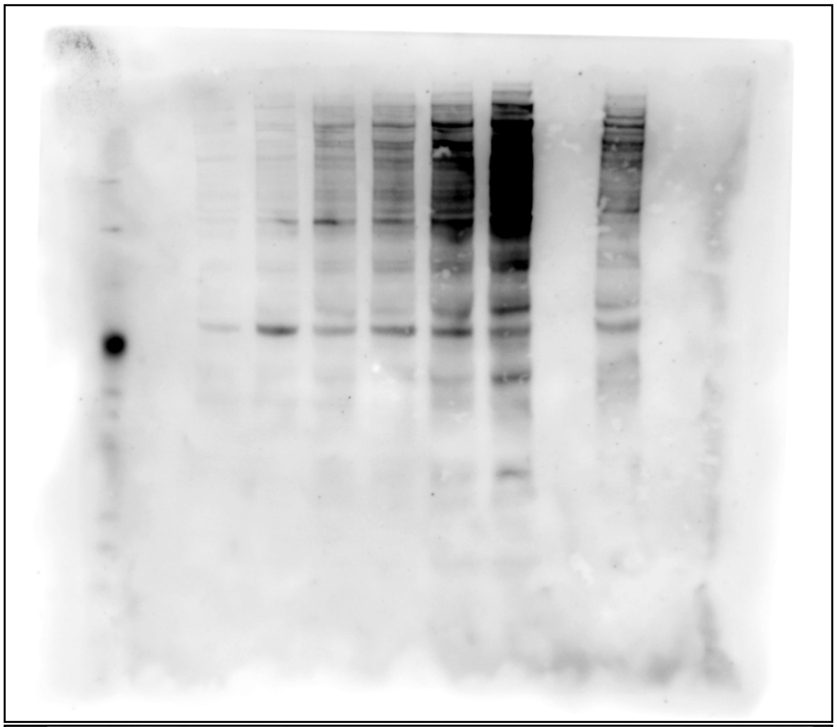

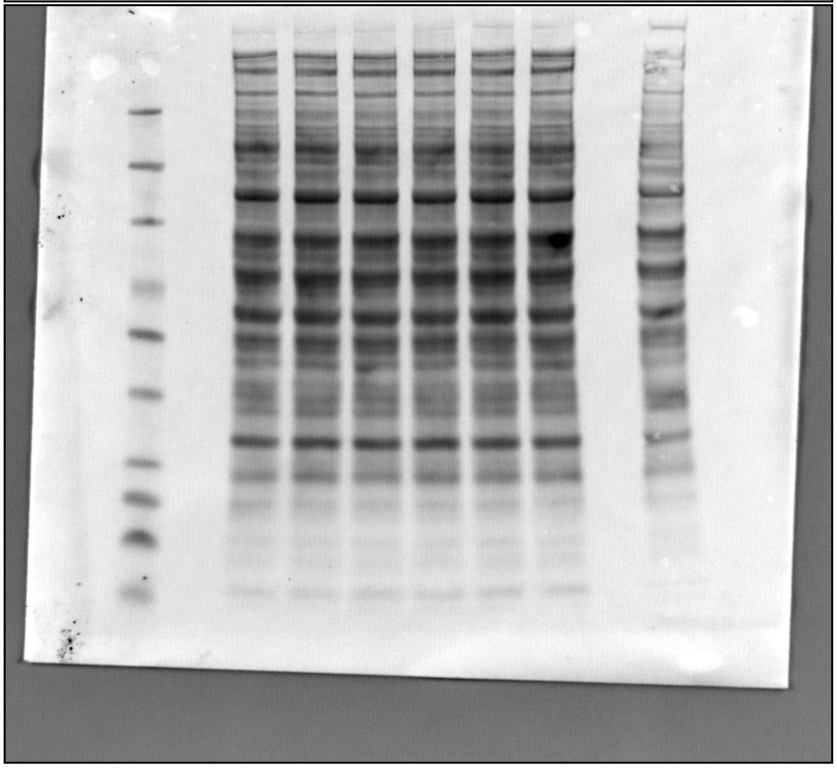

Full blot for Figure S3. Time-dependent cellular S-succination by maleate in UOK262 FH rescue (*FH+/+*) cells. Top: Chemiluminescence. Bottom: Ponceau.

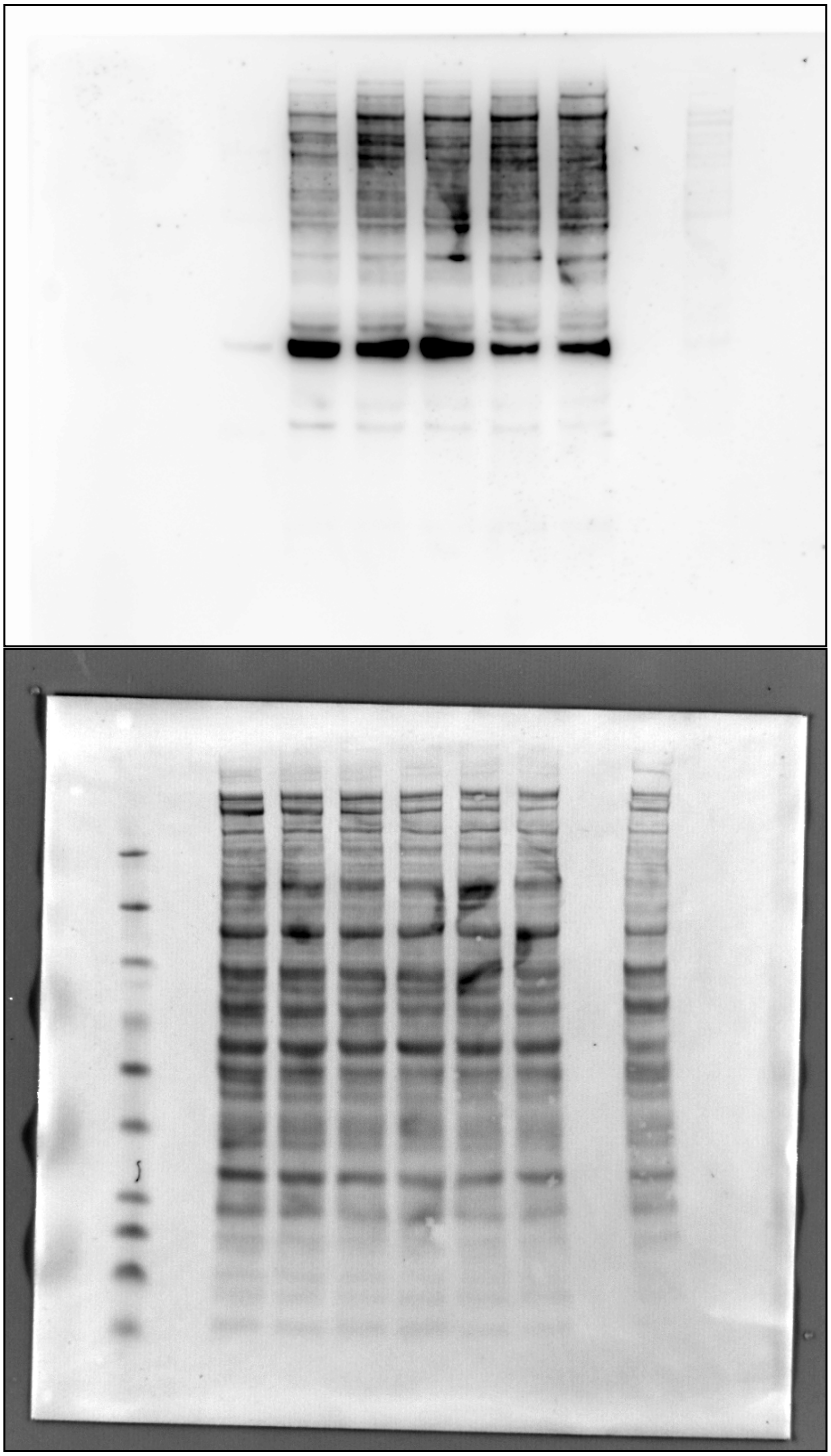

Full blot for Figure S3. Time-dependent cellular S-succination by 2-bromosuccinate in UOK262 FH rescue (*FH*+/+) cells. Top: Chemiluminescence. Bottom: Ponceau

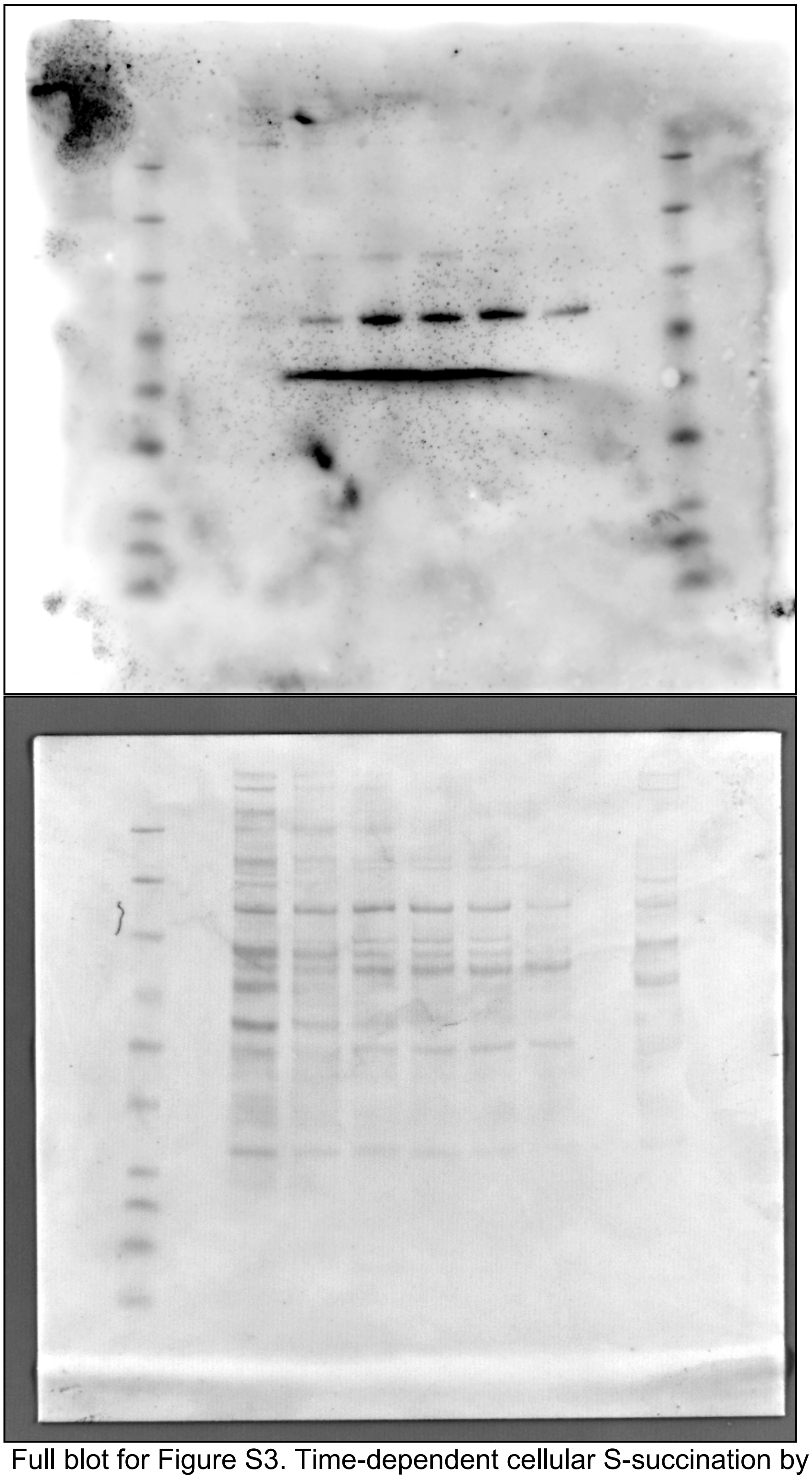

Full blot for Figure S3. Time-dependent cellular S-succination by dimethyl fumarate in UOK262 FH rescue (*FH+/+*) cells. Top: Chemiluminescence. Bottom: Ponceau

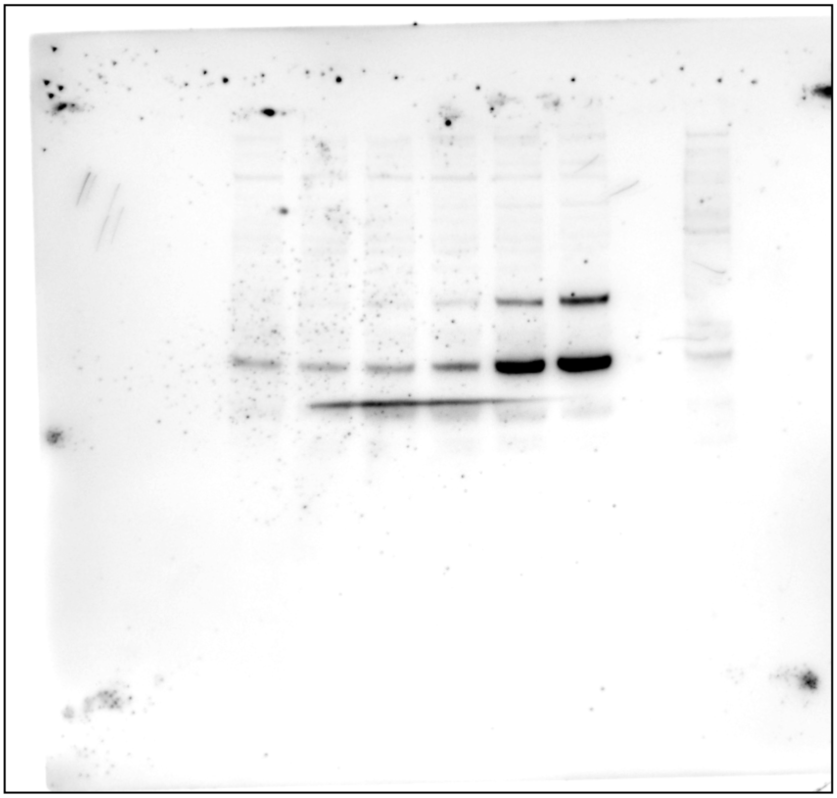

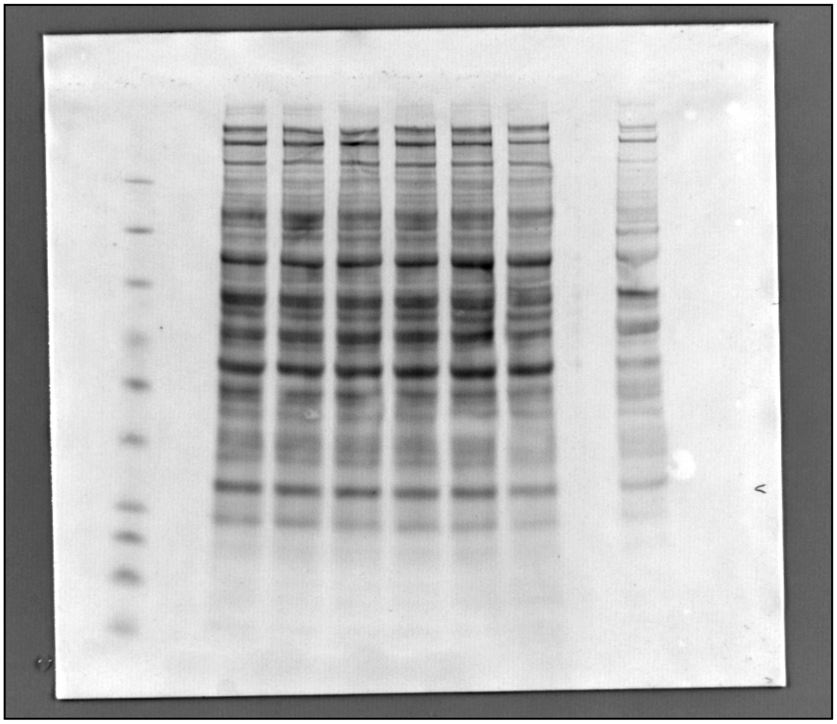

Full blot for Figure S3. Time-dependent cellular S-succination by ethyl fumarate in UOK262 FH rescue (*FH+/+*) cells. Top: Chemiluminescence. Bottom: Ponceau

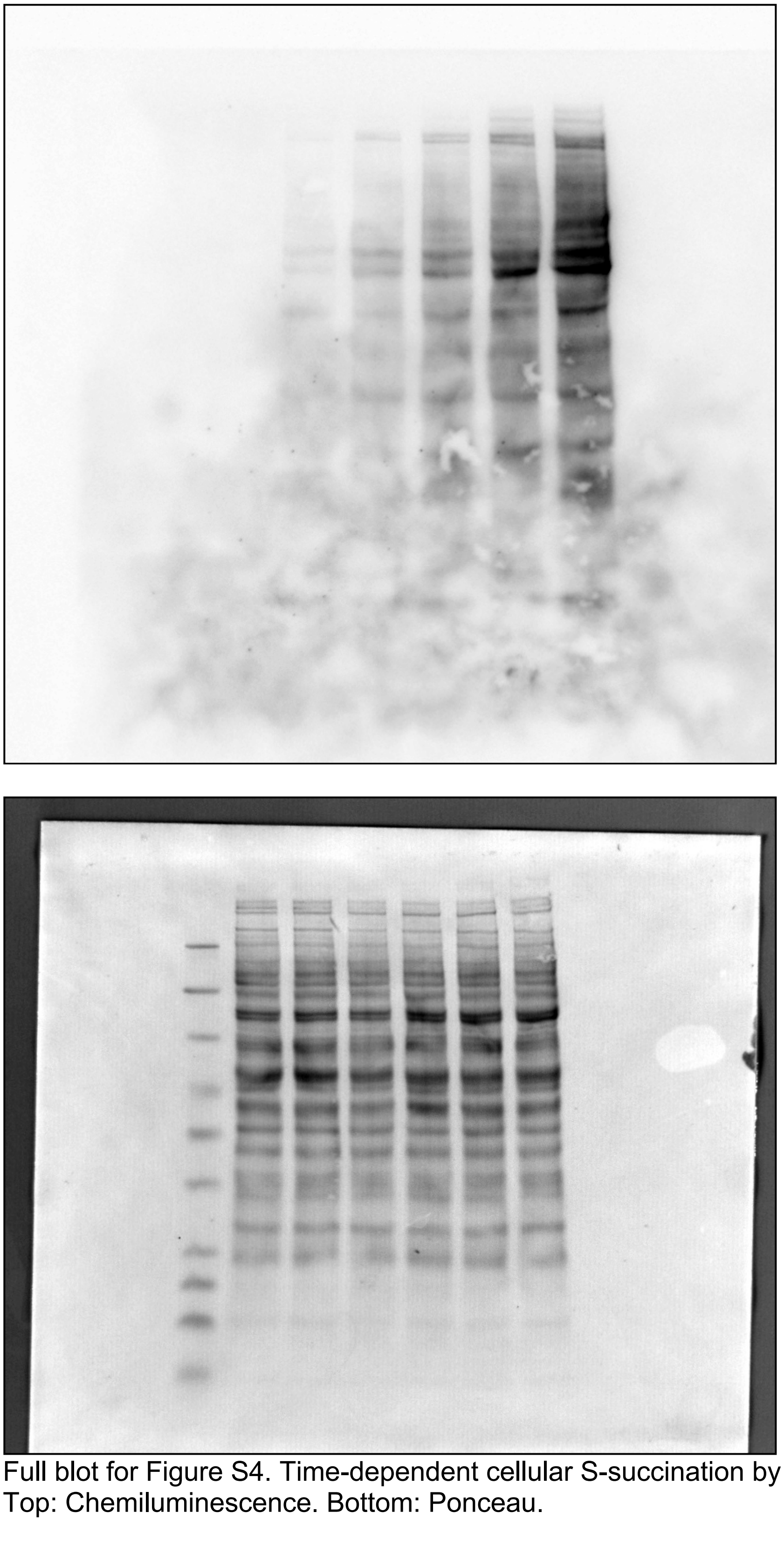

Full blot for Figure S4. Time-dependent cellular S-succination by maleate in HEK-293T cells. Top: Chemiluminescence. Bottom: Ponceau.

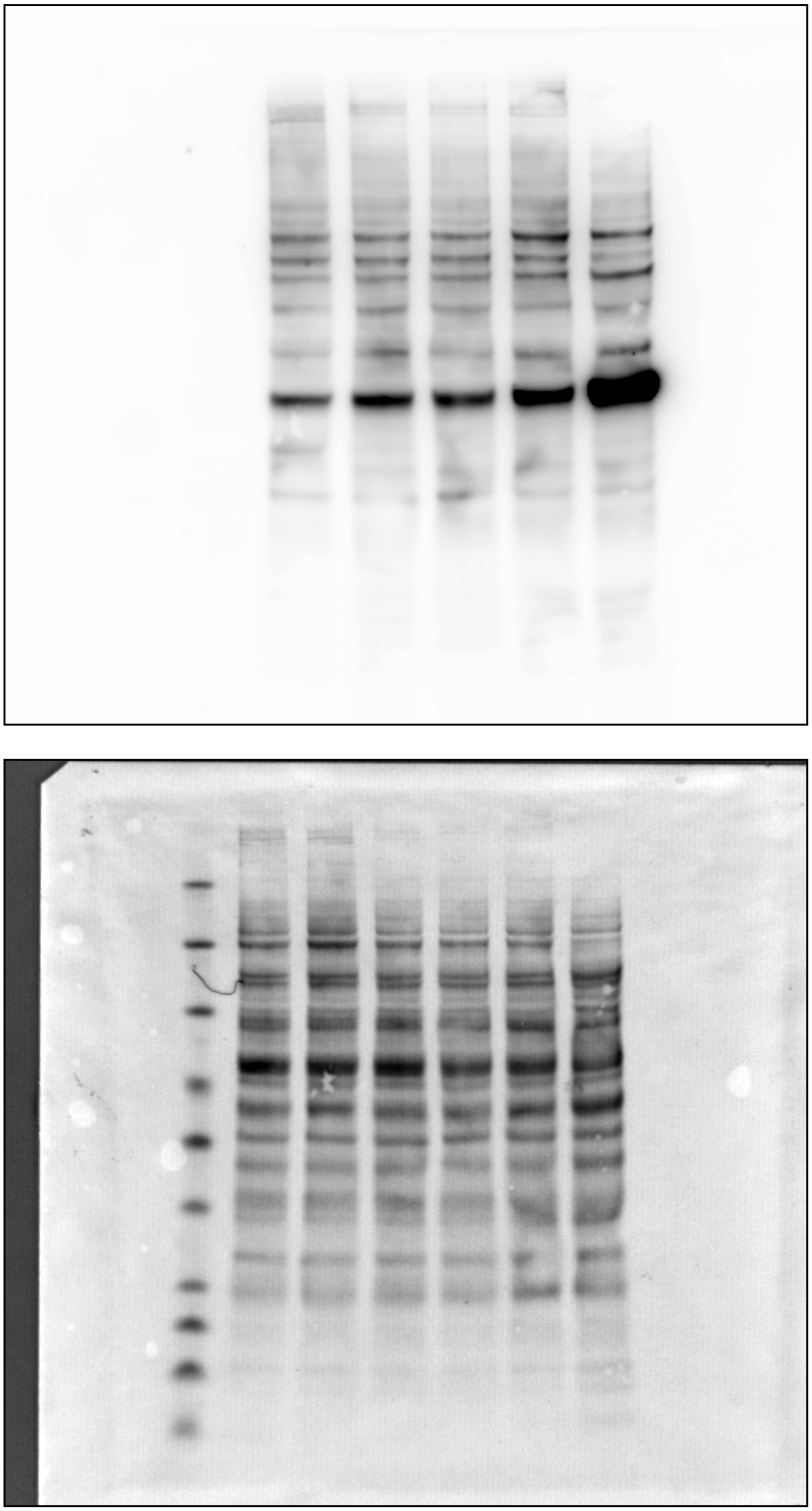

Full blot for Figure S4. Time-dependent cellular S-succination by 2-bromosuccinate in HEK-293T cells. Top: Chemiluminescence. Bottom: Ponceau.
